# Blaming Luck, Claiming Skill: Self-Attribution Bias in Error Assignment

**DOI:** 10.1101/2025.03.18.644058

**Authors:** Naoyuki Okamoto, Michael Taylor, Takatomi Kubo, Shin Ishii, Benedetto De Martino, Aurelio Cortese

## Abstract

Mistakes are valuable learning opportunities, yet in uncertain environments, whether a lack of reward is due to poor performance or bad luck can be hard to tell. To investigate how humans address this issue, we developed a visuomotor task where rewards depended on either skill or chance. Participants consistently displayed a self-attribution bias, crediting successes to their own ability while blaming failures on randomness, an effect that influenced their subsequent decisions. Computational modelling revealed two underlying mechanisms—a distorted perception of ability and a positivity bias in the skill condition. Notably, while distorted self-perception shaped behaviour, it did not affect confidence; instead, self-attribution bias led to overconfidence in external blame. These findings suggest a more complex picture in which self-attribution biases arise from both perceptual distortions and post-decision evaluations, highlighting the need for an interplay between experimental design and computational modelling to understand behavioural biases.

## Introduction

Using feedback (both positive and negative) to update beliefs and adjust behaviour lies at the heart of reinforcement learning (Doya, 2007; Sutton & Barto, 2018). This powerful learning strategy enables animals, including humans, to determine the correct course of action directly from their environment through trial and error. Algorithms based on RL have proven essential in driving the ongoing AI revolution (e.g., AlphaGo and AlphaFold) (Jumper et al., 2021; Silver et al., 2016). However, this learning strategy relies on the tacit assumption that humans (or artificial agents) can easily identify the source of an error leading to the lack of a reward. While this holds in most laboratory settings, where scenarios are designed with clear objectives and straightforward causes of errors, the complexity of the real world introduces multiple, often hidden, sources of errors. To learn efficiently, humans (and advanced AI systems) must discern whether a negative outcome stems from a genuine error or merely bad luck, such as randomness in the environment (Cortese, 2022; Rouault et al., 2019).

Here, we developed a novel visuomotor task to test how humans respond to errors caused by their performance or by factors independent of it. We found that, although participants can generally determine the source of an error, they tend to attribute the cause of a mistake to the environment while crediting themselves for success. This behavioural bias appears to manifest a general behavioural tendency well documented in psychology, known as self-attribution bias, often also called self-serving bias (Miller & Ross, 1975).

For example, in finance, self-attribution bias occurs when a trader attributes gains to their own skills but blames negative results on bad luck or other external factors (Chin et al., 2018; Hayes, 2023), leading to overconfidence and excessive risk-taking as traders overestimate their abilities (Hoffmann & Post, 2014). Similarly, in educational settings, students often credit high grades to their own intelligence and effort, whereas poor grades are blamed on unfair tests or inadequate students’ preparation while crediting successes to their own effective teaching practices (Covington, 1992; McAllister, 1996). Notably, self-attribution bias has been found consistently across different population strata, albeit with substantial variations due to age, cultural background and psychopathology (Mezulis et al., 2004). In the context of mental health, it has been linked to depression and the development of learned helplessness (Abramson et al., 1989; Seligman, 1972; Seligman & Maier, 1967). A recent study proposed self-attribution bias as a computational mechanism that might give rise to learned helplessness (Zamfir & Dayan, 2022).

However, most work on this phenomenon is limited to psychological traits (Shepperd et al., 2008) and their motivational underpinnings (Zuckerman, 1979). Only recently have a number of studies started to address how the valence of outcomes shapes learning in the context of reinforcement learning and Bayesian inference. While some authors have found that people harbour higher learning rates for positive outcomes (Garrett & Daw, 2020; Palminteri & Lebreton, 2022; Sharot & Garrett, 2016), others have shown the opposite effects (Gershman, 2015; Niv et al., 2012). To reconcile these findings, some have suggested the direction of valence-dependent learning asymmetries is due to beliefs about the causal structure of the environment (Dorfman et al., 2019).

In this study, we examined two competing mechanisms for self-attribution bias. One possibility is that the bias arises from post-hoc outcomes evaluation. On this positivity bias, individuals tend to give greater weight to positive feedback (successes) and downplay negative feedback (errors) in the condition in which they have control over the environment (i.e., in the skill condition). Alternatively, the bias may stem from an inflated internal representation of one’s abilities, which distorts how individuals perceive the environment.

To resolve this question, we incorporated several new key features into our experimental approach (Mancinelli et al., 2021; Zamfir & Dayan, 2022). First, we diverged from previous studies by employing a sensorimotor task rather than the more commonly used multi-armed bandit tasks, in which factors beyond mere choice play no role. This created a scenario in which the participant’s actual motor performance had a measurable effect on the score, with performance being continuously monitored prior to feedback. Second, we utilised a computational model incorporating a perceptual representation of one’s motor skills as an explicit threshold parameter. This model enabled us to test the effects of the alternative cognitive mechanisms described above. Finally, by measuring confidence on a trial-by-trial basis, we were able to examine the complex relationship between metacognition and the underlying cognitive strategy of credit and blame assignment.

## Results

### Task, error assignment and discovery of a novel self-attribution bias

Our new visuomotor task was inspired by the arcade game ‘whack-the-mole’ and modified to create scenarios in which rewards were sometimes tied to participants’ motor performance (‘skill’ condition) and sometimes independent (‘random’ condition). In each trial, participants had to quickly hit (within 800 ms) a cartoon mole that popped out of one of several ground holes shown on a touchscreen tablet. After hitting a mole, participants received a positive or negative score, which depended on the mole hit location in the ‘skill’ condition and was random in the ‘random’ condition. They then reported whether they believed this score was due to their action or mere randomness (see Fig. 1A, inference choice on the hidden task state ‘skill’ or ‘random’). Finally, participants reported their confidence about the accuracy of their decision (Fig. 1B). The task state (‘skill’ or ‘random’) was hidden and changed at random time points, unbeknownst to participants (see Supplementary Fig. 1 for sequence durations between switch points). Thus, participants had to weigh multiple sources of uncertainty to correctly infer the ongoing task state: uncertainty about the hidden rule and uncertainty about their motor movements in hitting the mole. Overall, participants (N = 51) inference accuracy was well above the theoretical chance level (mean accuracy 0.68 ± 0.08, chance level 0.5, Fig. 1C). While the task resulted in varying degrees of inference accuracy at the individual level, participants well adapted to the changing hidden state of the task (see two example participants in Fig. 1D). The task was calibrated such that participants would obtain positive scores with similar probabilities in both ‘random’ and ‘skill’ task states (supplementary Fig. 2). Overall, there was little difference between the two hidden states in participants’ inference rate of the actual condition (see Supplementary Table 1, Wilcoxon signed rank test on diagonal elements, i.e., true positive and true negative rates: Z = 1.87, *P* = 0.061).

**Figure 1:**
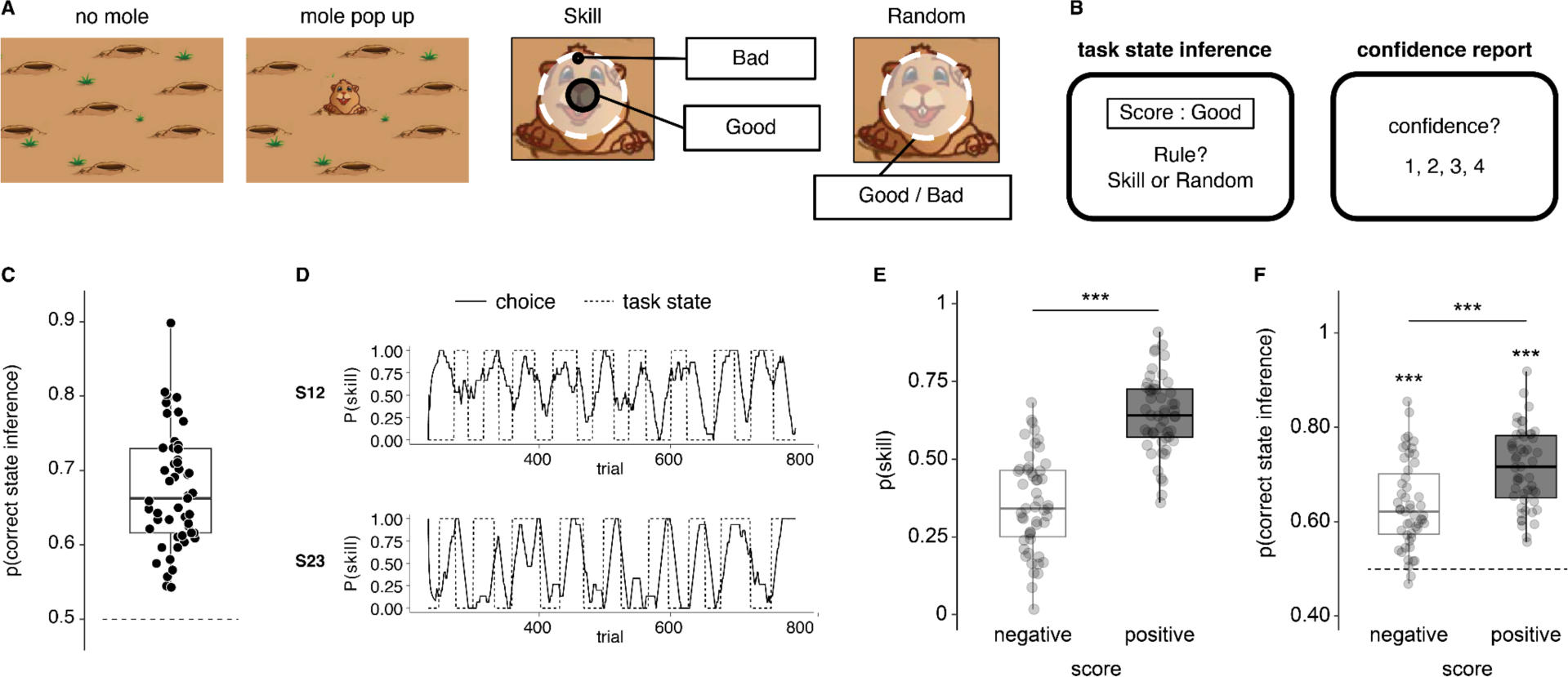
Task design, general behaviour results and self-attribution bias. (A) To score points in the task, participants had to hit (with a finger, using a touch screen) a mole that popped up and down quickly on a touchscreen. Following a target hit, participants saw the score. The task was divided into periods in which the score depended on the participant’s skill to hit the centre of the mole (skill condition) and periods in which the score was random (random condition). The length of each sequence, e.g., the number of trials between change points, followed a gamma distribution, with a mean duration of 28 trials. (B) After seeing the trial’s score, participants had to decide if they were in a skill or random condition. Next, participants reported their confidence in the correctness of their inference. (C) Participants’ overall accuracy in inferring the task state. (D) Example time courses of task state inference from two participants. Behaviour trajectories were smoothed with a backward time window of size n = 15 trials. (E) Participants’ probability of choosing the task state ‘skill’ as a function of the score obtained (‘negative’, ‘positive’). (F) Participants’ probability of correctly inferring the hidden task state as a function of the score obtained (‘negative’, ‘positive’). In panels C, E and F, dots represent individual participants, and box plots the median and interquartile ranges. N = 51 human participants, *** *P* < 0.001 (Wilcoxon signed-rank test).

Participants displayed a clear self-attribution bias. In short, they chose the ‘skill’ state significantly more often when they received a positive score following a mole hit than when they received a negative score (Z = 5.99, *P* < 0.001, Fig. 1E). In turn, participants’ probability of making a correct state inference also differed depending on the score received, with overall higher accuracy in trials whose score was positive compared with trials whose score was negative (Z = 5.22, *P* < 0.001, Fig. 1F). Note that inference accuracy was significantly above chance in both cases (test for inference accuracy > 0.5; negative scores: Z = 6.14, *P_FDR_* < 0.001, positive scores: Z = 6.21, *P_FDR_* < 0.001). Similarly, participants had slower choice reaction times following negative vs positive scores (Supplementary Fig. 3A).

Higher inference accuracy following positive scores could have reflected a nonspecific task confound, making positive scores easier to evaluate. To check for this, we first tested the effect of the score (negative vs positive), the true state of the task (skill vs random) and their interaction on participants’ inference accuracy using a two-way repeated measures ANOVA (Fig. 2A). This analysis revealed a significant interaction (F_(1, 50)_ = 127.3, *P* < 0.001), as well as a significant main effect of score (F_(1, 50)_ = 60.2, *P* < 0.001), while the effect of task state was not significant (F_(1, 50)_ = 0.17, *P* = 0.68). Post-hoc pairwise comparisons showed drastic differences in inference ability, with higher accuracy following negative compared to positive scores in the ‘random’ state (Z = −5.01, *P_FDR_* < 0.001) but a mirrored effect of higher accuracy following positive compared to negative scores in the ‘skill’ state (Z = 6.14, *P_FDR_* < 0.001). Accordingly, negative scores led to higher accuracy in the ‘random’ state compared to the ‘skill’ state (Z = −4.86, *P_FDR_* < 0.001), while positive scores led to higher accuracy in the ‘skill’ state compared to the ‘random’ state (Z = 5.56, *P_FDR_* < 0.001).

**Figure 2:**
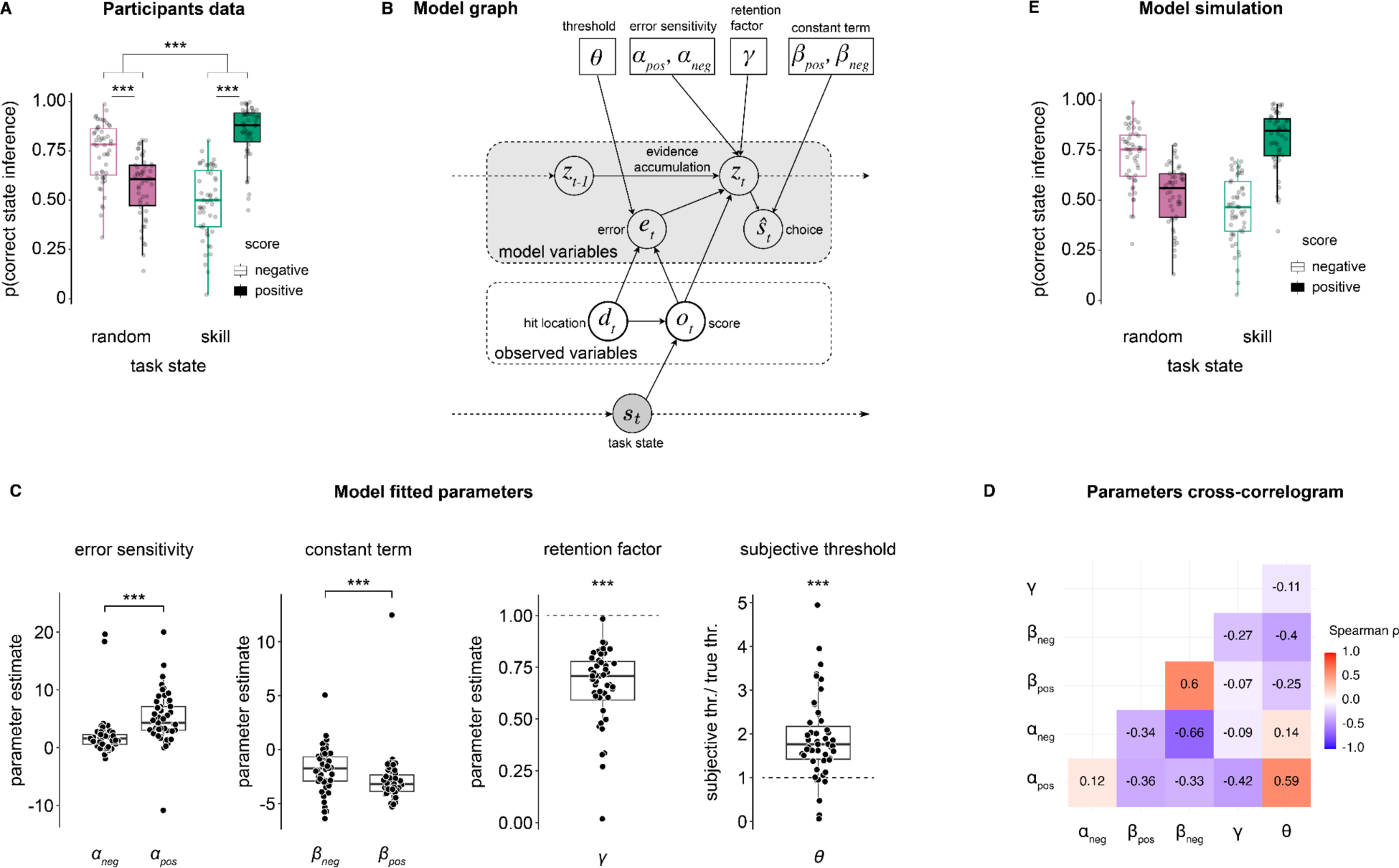
Inference accuracy and computational model. (A) Participants’ probability of correctly inferring the hidden task state as a function of the score obtained (‘negative’, ‘positive’) and the true task state (‘random’, ‘skill’). The effects of score and task state and their interaction on participants’ inference accuracy were statistically evaluated with repeated measures two-way ANOVA. Dots represent individual participants, boxplots the median and first/third quartiles, and whiskers the minimum and maximum of the data range. (B) Graph illustration of the computational model. The model features a set of observable variables [the distance of the hit location from the mole centre *d_t_* and the score obtained *o_tt_*] and latent variables (evidence accumulation *z_t_* and the error *e_t_*). The model’s output is the estimated state (choice) 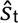. The key parameters that control the model’s behaviour are the agent’s subjective threshold between the centre and edge of the mole (*θ*), the error sensitivity modulating the effect of the error (score-dependent, *α*_pos_, *α*_neg_), the retention factor modulating the evidence accumulation (*γ*), and a constant term modulating the decision boundary (separately for positive and negative outcomes, *β*_pos_, *β*_neg_). (C) Model parameters are fitted through negative log-likelihood minimisation. The four panels show, from left to right, the score-dependent error sensitivity *α*, the score-dependent constant term *β*, the retention factor *γ*, and the ratio between the subjective threshold *θ* and the true threshold. (D) Parameters cross-correlogram. We used Spearman rank correlation across all parameters’ distributions (each parameter was fitted at the individual participant level) to compute the cross-correlogram. (E) Model simulations of p(correct state inference). Simulations based on the estimated parameters and the original trial time courses. Each dot represents one participant/agent for all scatter plots, *** *P* < 0.001.

We then tested whether participants’ mole-hitting patterns provided additional insight into their behavioural strategy. To do this, we analysed the hit locations, precisely the distance from the mole centre, in three ways: (i) statically, (ii) over time and (iii) in relation to previous scores and decisions. First, we confirmed that hits were overall closer to the centre when they resulted in positive compared to negative scores (as expected given the task design, Supplementary Fig. 3B). Interestingly, we found that where participants hit the moles in relation to the centre differed depending on the objective task state (see Supplementary Fig. 4A-B, hits were closer to the centre in the skill state than the random state). Furthermore, on a more granular level, obtained score (negative, positive) and task state inference (skill, random) impacted where participants hit the mole on the following trial (Supplementary Fig. 4C). Participants improved their hit precision following a negative score but decreased it following positive scores. This result held both when participants reported ‘random’ or ‘skill’ but with a more substantial effect when the choice was ‘skill’ (main effect of score F_(1, 50)_ = 159.68, *P* < 0.001; main effect of inference choice F_(1, 50)_ = 24.96, *P* < 0.001; interaction between inference choice and score: F_(1, 50)_ = 71.85, *P* < 0.001). The magnitude of the difference was larger after a positive score compared to a negative score, whichever rule they chose (random: Z = −5.32, *P_FDR_* < 0.001; skill: Z = −6.21, *P_FDR_* < 0.001). Together, these results suggest that participants dynamically adjusted their hit location. However, there was no evidence that they did so strategically, in which case we should have seen an effect only in the ‘skill’ state and not in both, as reported here.

### Behavioural signatures of self-attribution bias and computational modelling

How does self-attribution bias lead to incorrect error attribution, and what are its computational underpinnings? More specifically, at what stage does this error attribution occur? To address these questions, we tested two alternative cognitive mechanisms.

The first mechanism operates at the feedback level. In this scenario, much like positivity or confirmation bias (Palminteri & Lebreton, 2022; Sharot & Garrett, 2016), participants tend to neglect negative feedback by misattributing it to external factors. The second, more pernicious mechanism involves an inflated estimation of one’s abilities (motor in this task); as a result, participants are less inclined to attribute errors to themselves.

To test these two possibilities, we developed a computational model of our task in which scores derive from the joint effect of the hit location (i.e., distance from the centre of the mole) and the hidden task state (Fig. 2B). The model features a leaky error-evidence accumulator updated on every trial by an error term computed from the mismatch between the expected hit location (centre, edge) and the score obtained (positive, negative). The key parameter modelling participants’ perception of their own motor ability is *θ*, which determines the subjective threshold between the centre and the edge of the mole, i.e. the boundary coding a switch in score from positive to negative in the ‘skill’ state. Additional parameters include score-dependent *α*, acting as an error sensitivity that modulates the strength of the error update on the evidence accumulator; *γ*, a retention factor modulating the accumulation process; and a score-dependent constant term *β* modulating the decision variable itself (see methods for full model description).

We first fit the model’s free parameters to participants’ behaviour data through a log-likelihood minimisation procedure. Parameters’ fits (Fig. 2C-D) revealed several interesting behaviour features. First, the error sensitivity *α* was larger for positive scores compared to negative scores (Z = 5.08, *P* < 0.001), indicating that participants weighted more feedback that resulted from positive scores (in line with work on confirmation bias in reinforcement learning (Palminteri, Lefebvre, et al., 2017; Palminteri & Lebreton, 2022)). Similarly, the constant term *β* was larger (more negative) for positive scores compared to negative scores (Z = −4.07, *P* < 0.001). Note that the constant term regulates the decision boundary in reporting ‘skill’ and ‘random’, and in our model, a more negative constant term shifts the decision boundary towards ‘skill’ inference choice. Thus, the parameter fitting highlighted the overall tendency for positive scores to lead to skill choices. Third, the error evidence accumulation process was leaky rather than lossless, as the retention factor *γ* was significantly smaller than 1 (Z = −6.21, *P* < 0.001), reflecting the noisy nature of the participants’ error evaluation process. Fourth, and most critical to test one of our hypotheses, the ratio of the subjective threshold *θ* and the true threshold was significantly larger than 1 (Z = 5.48, *P* < 0.001), indicating that participants consistently overestimated their motor skills by liberally assessing the mole central hit zone (giving positive scores in the ‘skill’ hidden state). Participants’ bias in assigning positive scores to their own ability resulted from a distortion of their perceived ability, in this case, motor ability.

Comparisons between the full model and alternative, simpler versions (e.g., with a single score-independent error sensitivity or constant term, true threshold, and no retention factor) confirmed that the full model better accounted for participants’ choice strategies (see model comparison results in Supplementary Fig. 5). In addition, the model comparison further highlighted that the perceptual inflation (subjective threshold *θ*) played a larger role than overall positivity bias (error sensitivity *α* and constant term *β*) in determining behaviour, given the larger AIC difference from the full model when this parameter was fixed to the true threshold value (Supplementary Fig. 5).

Next, we simulated new choice data using the parameters obtained from the fitted full model. We verified that the model could accurately capture the key behavioural signature of self-attribution bias in error assignment (Fig 2E, see also Supplementary Fig. 6). These results were extended with a set of simulations with pre-set parameter values (e.g., fixing the subjective threshold *θ* to the true value, fixing *α* and *β* to be equal for both positive and negative scores), revealing that under different parameter regimes, behaviour changed drastically. Under these circumstances, the model failed to replicate participants’ inference accuracy patterns (Supplementary Fig. 7). In particular, simply fixing *θ* to the true value cancelled a significant portion of the self-attribution bias (absence of bias in the ‘random’ state and minimised bias in the ‘skill’ state, Supplementary Fig. 7B). The oracle model (true threshold, single error sensitivity and single constant term) displayed high overall inference accuracy across all task states and scores without biases, as expected (Supplementary Fig. 7H).

Finally, we validated the model fitting procedure by performing a parameter recovery analysis with simulated data. Note that although some correlations between parameters were relatively high (Fig. 2D), we were able to individually recover all parameters with good reliability in the parameter recovery analysis (Supplementary Fig. 8). Results thus confirmed the model’s robustness, with correlations between fitted parameters and recovered parameters at *r* ≥ 0.74.

### Self-Attribution Bias Determines Future Behaviour Strategy

Participants adapted smoothly to hidden switches in the task state (from ‘random’ to ‘skill’ or vice versa). As shown in Fig. 3A, the probability of making a correct inference dropped below 0.4 immediately after a task state switch, then recovered and stabilised over the following 7–8 trials. Because we observed that inference accuracy varied with the hidden task state (i.e., Fig. 2A), we first asked whether the direction of the switch (‘random’ → ‘skill’ vs. ‘skill’ → ‘random’) differentially influenced participants’ inference accuracy. We specifically focused our statistical analysis on trials following a switch (in Fig. 3A, trials 1-8). A repeated-measures ANOVA revealed no significant difference in inference accuracy between the two switch directions (F_(1, 50)_ = 1.70, *P* = 0.20), beyond the expected general increase in performance over trials from switch time (main effect of time: F_(1, 50)_ = 66.6, *P* < 0.001) and no significant interaction (F_(1, 50)_ = 1.84, *P* = 0.079). None of the pairwise comparisons between the two trajectories resulted in significant differences (all *P*_FDR_ > 0.1). Notably, model simulations closely captured this behavioural time course (Fig. 3B, main effect of time, no effect of switch direction, no significant interaction, and all pairwise comparisons *P*_FDR_ > 0.19). See Supplementary Fig. 9 for simulations with different parameter regimes.

**Figure 3:**
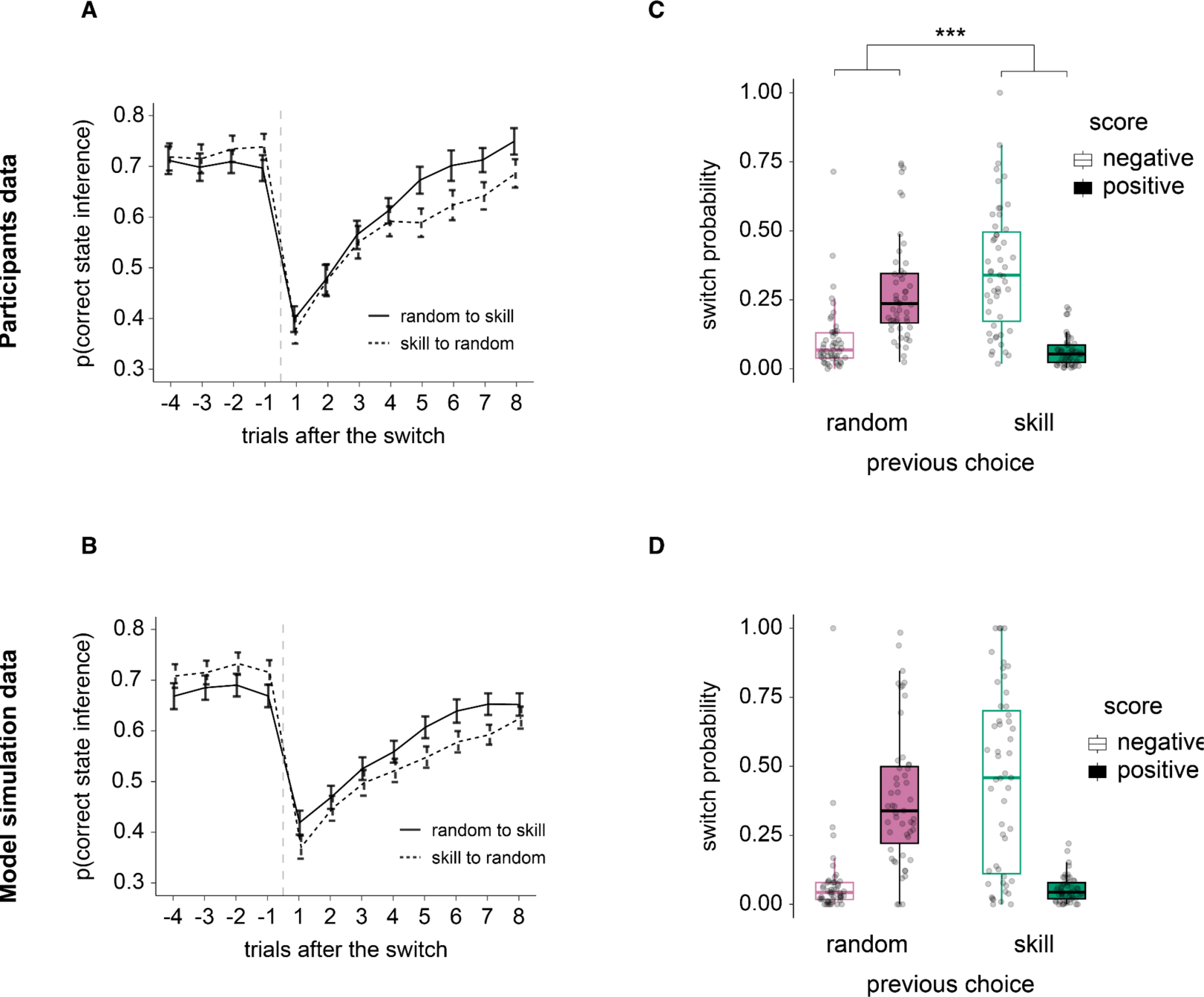
Inference choices around task change points and effect on subjective switch probability. (A) Participants’ probability of correctly inferring the hidden task state as a function of the trial from the objective task state switch and the direction of the switch (solid line ‘random → skill’, dotted line ‘skill → random’). The main effects of trial and switch direction and their interaction on participants’ inference accuracy were statistically evaluated with repeated measures ANOVA. Solid/dotted lines represent the group average, and the error bars represent the standard error of the mean. N = 51 participants. (B) Same as in A, but with artificial data generated from a simulation. N = 51 simulated agents. (C) Participants’ probability of switching their choice (e.g., from random to skill or skill to random) as a function of the score obtained (‘negative’, ‘positive’) and their choice in the previous trial (‘random’, ‘skill’). The main effects of score and choice and their interaction on participants’ switching probability were statistically evaluated with repeated measures two-way ANOVA. Dots represent individual participants, boxplots the median and first/third quartiles, and whiskers the minimum and maximum of the data range. (D) Same as in C, but with artificial data generated from a simulation. For all plots, N = 51 participants (A, C) or simulated agents (B, D); *** *P* < 0.001.

We examined how self-attribution bias might shape participants’ subsequent choices. Because the hidden task state (‘random’ vs. ‘skill’) and the score (positive vs. negative) jointly influenced correct inferences, we hypothesised that belief about the task state (previous choice at trial *t-1*) and the current score (at trial *t*) would together determine whether participants switched their choice at trial *t* (e.g., from ‘skill’ to ‘random’, or vice versa, Fig. 3C). Indeed, there was a strong interaction between current score and previous choice on the probability of switching (F_(1, 50)_ = 75.43, *P* < 0.001). We also found a main effect of the score (F_(1, 50)_ = 15.06, *P* < 0.001) but no effect of the previous choice alone (F_(1, 50)_ = 1.57, *P* = 0.22). Specifically, if participants had reported ‘random’ in the previous trial, receiving a negative score on the current trial led to a significantly lower chance of switching than receiving a positive score (Z = −5.11, *P_FDR_* < 0.001). This pattern reversed if participants had previously chosen ‘skill’: after a positive score, the probability of switching was lowest (Z = 5.81, *P_FDR_* < 0.001). In other words, participants switched more often if they believed the hidden state was skill but received a negative score or if they believed it was random and received a positive score. These findings demonstrate how self-attribution bias—attributing successes to one’s own skill and failures to external randomness—directly shapes behaviour and decision strategies. Importantly, our model captured this strategy-updating pattern well (Fig. 3D).

### Confidence and its dissociation from inference accuracy

During the task, participants also reported confidence judgments about the correctness of their inferential choices. Using a repeated measures two-way ANOVA, we confirmed a significant main effect of choice correctness (F_(1,50)_ = 176.5, *P* < 0.001) and of score valence (F_(1,50)_ = 39.69, *P* < 0.001) on confidence, but no interaction (F_(1,50)_ = 1.22, *P* = 0.27). Decision confidence in correct trials was higher than in error trials (Fig 4A, Z = 5.01, *P* < 0.001). In line with previous findings that confidence also reflects valence (Kaanders et al., 2022; Lebreton et al., 2019), participants’ confidence was highest after receiving positive scores (Z = 5.01, *P* < 0.001, fig. 4B). Thus, self-attribution bias also appears to impact the subjective evaluation of one’s actions.

**Figure 4:**
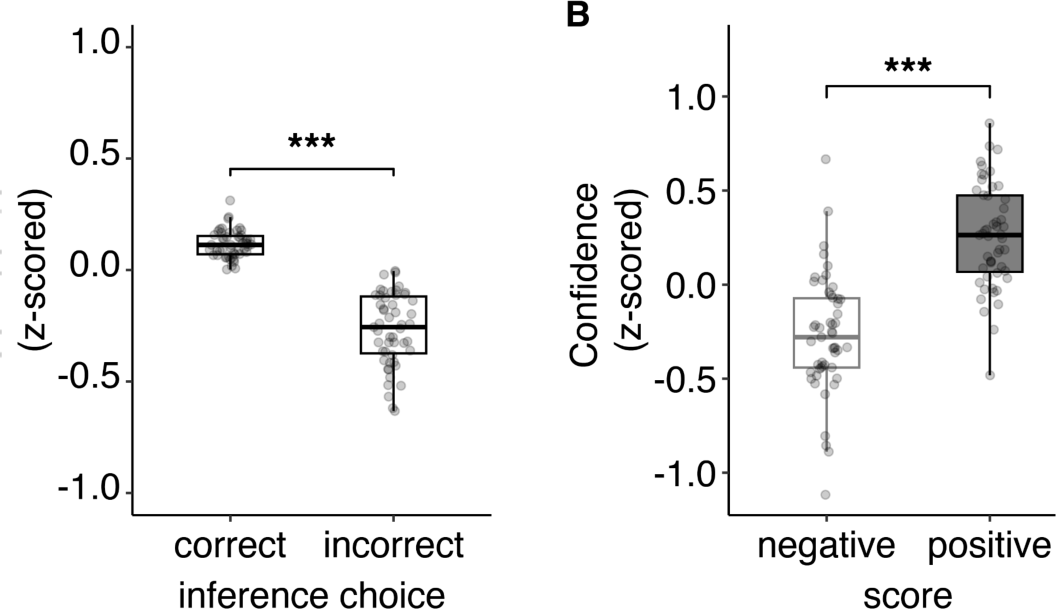
Confidence about inference correctness. (A) Participants’ confidence about the task states inference. Data plotted separately for correct and error trials, highlighting the typical signature of decision confidence, with higher confidence in correct trials compared to error trials. (B) Confidence as a function of the score obtained (‘negative’, ‘positive’). As with task state inference accuracy, confidence was higher following positive scores. Dots represent individual participants, boxplots the median and first/third quartiles, and whiskers the minimum and maximum of the data range. Confidence was z-scored within participants. *** *P* < 0.001

As done previously with inference accuracy, we tested whether scores (negative vs positive) and hidden task state (‘skill’ vs ‘random’) might have differentially impacted confidence judgements (two-way repeated measures ANOVA, Fig. 5A). Interestingly, confidence displayed a different picture, with a strong main effect of score (F_(1,50)_ = 44.75, *P* < 0.001, i.e., participants reported higher confidence in positive than negative scores: ‘random’ state: Z = 4.11, *P_FDR_* < 0.001; ‘skill’ state: Z = 5.39, *P_FDR_* < 0.001), a main effect of task state (F_(1,50)_ = 16.68, *P* < 0.001), and a significant interaction (F_(1, 50)_ = 10.1, *P* = 0.0025). This interaction reflected the main effect of the task state on confidence in the presence of negative scores, in which confidence was higher in the ‘random’ than ‘skill’ state (Z = −4.99, *P_FDR_* < 0.001, simple effect) but not of positive scores, in which confidence was similar in both task states (Z = −0.56, *P_FDR_* = 0.58, simple effect).

**Figure 5:**
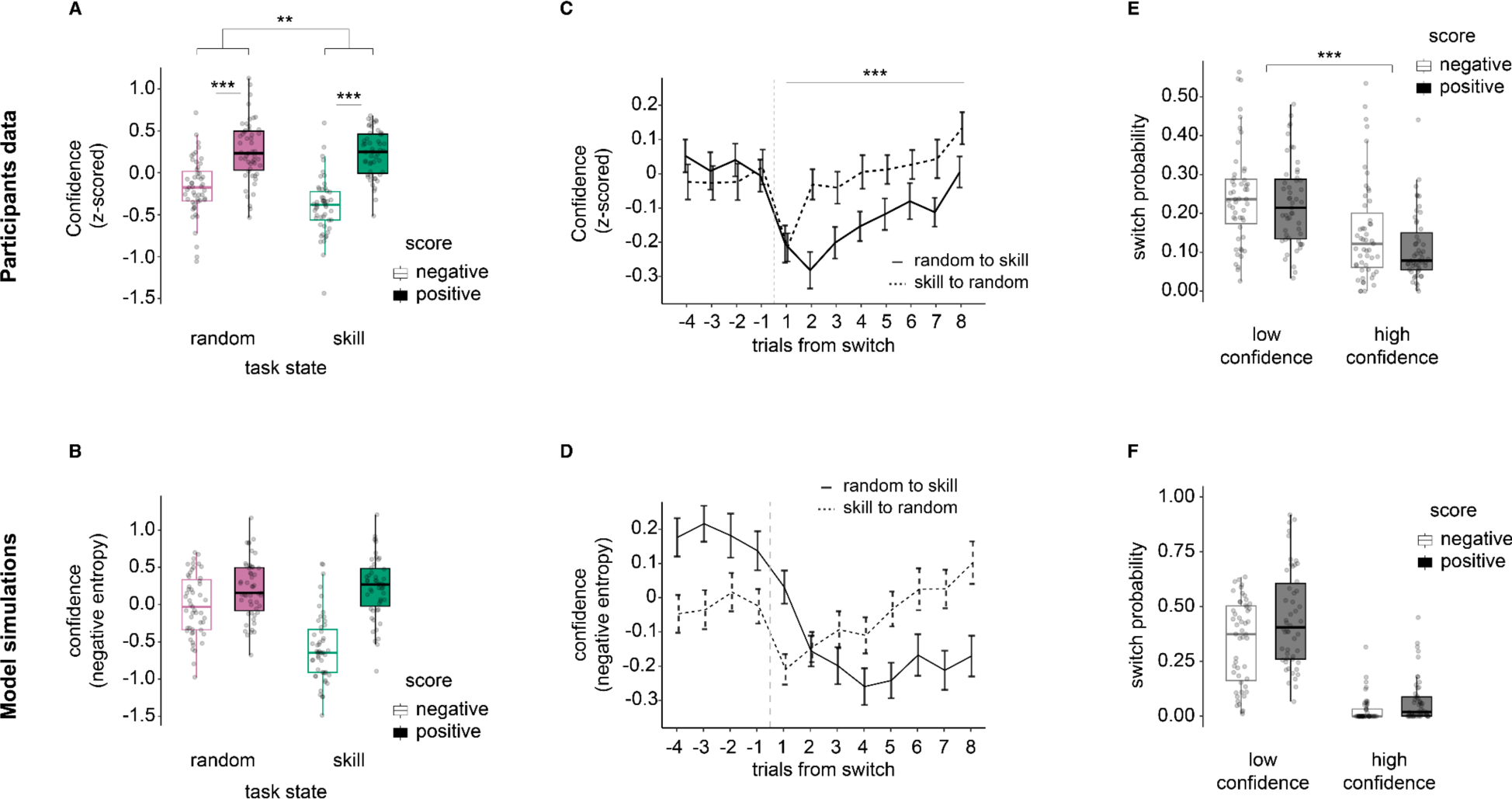
Confidence about the correctness of the inference. (A) Participants’ confidence about their hidden task state inference as a function of the score obtained (‘negative’, ‘positive’) and the true task state (‘skill’, ‘random’). The main effects of score and task state and their interaction on participants’ inference accuracy were evaluated with repeated two-way ANOVA. For all plots, dots represent individual participants, and box plots represent the median and interquartile ranges. (B) Same as in A, but with artificial data generated from a simulation. Note that the model was only optimised using inference decision data, not confidence. However, since the model decision module was based on a logistic function, we used the z-scored negative entropy about the model’s hidden task state inference decision as a proxy for confidence. (C) Time series of participants’ confidence around the switch (e.g., from random to skill or skill to random). The solid line is the mean of mean confidence within participants, and the bar is the standard error of the mean. (D) Same as in C, but with artificial data generated from a simulation. (E) Switch probability of the inference as a function of the score obtained (‘positive’, ‘negative’) and the confidence (binarised with its mean within participants). The main effects of score and confidence and their interaction on switch probability of the inference were evaluated with repeated two-way ANOVA. (F) Same as in E, but with artificial data generated from a simulation. ** *P* < 0.01, *** *P* < 0.001

Note that we do not have access to model-based confidence judgements for the computational model since it was fit and optimised simply on the inference decision data, not confidence. However, as a straightforward proxy of confidence for the model-based simulations, we used negative entropy (entropy represents the uncertainty in the model-based decision; see Supplementary Fig. 10 confirming the correspondence between participants’ confidence judgements and the model’s negative entropy). Even though its primary purpose was not to model confidence, our computational model still closely replicated the confidence pattern found in participants (Fig. 5B, see also Supplementary Fig. 11). Further simulations with different parameter regimes reinforced the qualitative differences between inference accuracy and confidence uncovered earlier (Supplementary Fig. 12, persistent valence effect on confidence independent of task state).

This valence effect on confidence suggests a potential metacognitive dissociation between choice accuracy and confidence (e.g., comparing Fig. 2A vs. 5A) that depended on the task state-score condition. Overall meta-d’ (a measure of how well confidence tracks decision accuracy (Maniscalco & Lau, 2012)) did not differ across task states (Z = 0.056, *P* = 0.96) nor score (Z = 0.58, *P* = 0.57, see Supplementary Fig. 13A-B). Interestingly however, participants displayed a specific drop in metacognitive ability in trials with negative scores in the skill state (see Supplementary Fig. 13C-D). Self-attribution bias thus appeared to make participants overconfident not in attributing successes to themselves but in attributing blame to external causes (i.e., negative scores blamed on environmental randomness).

On a trial-by-trial level, confidence reflected changes in task uncertainty, with participants giving their lowest ratings on average in the trials immediately following a change in the hidden task states (Fig. 5C). However, confidence judgements differed starkly from the state inference accuracy trajectories around these time points (i.e., Fig. 3A). The pattern of confidence judgements was asymmetric with respect to the direction of the transition (random → skill, skill → random), with a significant main effect of switch direction (F_(1, 50)_ = 12.7, *P* < 0.001), besides a general main effect of trials post-switch (F_(1, 50)_ = 8.19, *P* < 0.001), and a lack of interaction (F_(1, 50)_ = 1.49, *P* = 0.17). In short, reported confidence decreased when the hidden state changed, suggesting that participants generally noticed task environment changes, even without direct, explicit feedback about the task state or their choices. Nevertheless, confidence recovered faster when the hidden state switched from skill to random than when it switched from random to skill. Although exaggerated, the computational model displayed a similar difference, with a slow confidence recovery rate following a random → skill task state transition (Fig. 5D). Note that plotting a longer time-scale (from −16 trials pre-switch to +15 trials post-switch) shows that both participants and model had similar tendencies: overall higher confidence in the random condition (Supplementary Fig. 14). Simulations done with specific parameter settings revealed one more important piece of the puzzle (Supplementary Fig. 15). All simulations in which the subjective threshold parameter *θ* was fixed to the true value displayed confidence patterns that were very similar to participants’ own confidence ratings (Supplementary Fig. 15B, D, F, H). This surprising result thus reveals a dissociation between confidence and choice, in which choice is affected by the perceptual distortion of self-attribution bias, but confidence is not.

In line with previous work (Folke et al., 2016), we finally sought to verify whether confidence informs future choices. Participants had a higher probability of switching when they reported low confidence (main effect of confidence F_(1, 50)_ = 73.70, *P* < 0.001) but also when the score obtained was negative (main effect of score F_(1, 50)_ = 6.25, *P* = 0.016, no significant interaction F_(1, 50)_ = 1.50, *P* = 0.28, Fig. 5 E). The model captured these effects. These results are reminiscent of previous findings suggesting confidence regulates behaviour updates (Nassar et al., 2010; Vaghi et al., 2017) but also further highlight the strength of valence distortions in controlling behaviour.

## Discussion

This study aimed to understand how people attribute or misattribute negative feedback to their own perceptual errors or from environmental noise (i.e., bad luck). We compared two main hypotheses regarding the origin of the self-attribution bias we isolated in the context of a controlled visuomotor task. The first hypothesis considers that bias arises from a self-confirmatory mechanism, similar to positivity or confirmation biases (Kaanders et al., 2022; Palminteri & Lebreton, 2022; Tarantola et al., 2021). In this view, individuals outweigh positive feedback (successes) and underweight negative feedback (errors) at the moment of the feedback evaluation. Instead, the alternative hypothesis states that the bias results from a systematic misestimation of one’s own abilities—in this case, motor ability—leading individuals to attribute errors to external factors due to an inflated perception of personal competence. Importantly, each explanation implies fundamentally different underlying cognitive processes, necessitating distinct interventions if one aims to develop a behavioural nudge to reduce the bias.

Using behavioural and computational modelling, our study provides clear evidence that a self-confirmation bias can emerge from a systematic inflation of one’s perceived ability. We show that the parameter *θ* that controls the perceptual threshold is greater than the true threshold (on average, almost twice as large), indicating that participants consistently overestimated their motor skills by liberally considering the mole central hit zone (giving positive scores in the skill hidden state). A result confirmed by our simulations and model comparison analyses. This pattern of results implies that self-attribution bias appears to be an intrinsic component of basic sensory processing related to the self, influencing behaviour directly from the outset of the action, beyond a post-hoc rationalisation for negative feedback. There is growing evidence for early modulation of perceptual processes by a variety of contextual constraints such as environment, task and even cognitive demands (Banerjee et al., 2020; Kelly et al., 2020; Schaffner et al., 2023).

However, our model-based analysis has portrayed a more complex picture in which the overestimation of the skill (while key to triggering the bias) is not the only process that plays a role in generating the effect we report. We also show that, in line with work on confirmation bias in the context of reinforcement learning (Palminteri, Lefebvre, et al., 2017; Palminteri & Lebreton, 2022), participants weighted more feedback that resulted from positive scores, in line with our first hypothesis. Intriguingly, the confirmation (or positivity) bias might even act as a compensatory mechanism that enhances behavioural performance under the presence of self-attributional perceptual distortions (e.g., see Supplementary Fig. 7C for simulation results with an agent harbouring perceptual inflation of ability but no confirmation/positivity biases in error sensitivity and decision parameters, showing reduced inference accuracy in the skill condition). This speculation was supported by a second preliminary piece of evidence, in that the extent of the perceptual distortion in one’s ability correlated with the strength of the positivity/confirmation bias (see Supplementary Fig. 16). Nevertheless, while intriguing, these results are exploratory, and future work with carefully designed behaviour task, modelling and simulations is necessary.

A recent study on self-attribution bias revealed dynamic, reciprocal relationships between attributions and self-beliefs, supporting the attribution-self-representation cycle theory (Zamfir & Dayan, 2022). They show that participants were more likely to update their beliefs about their abilities when they attribute outcomes to themselves rather than external factors. This aligns with our findings, which show a higher error sensitivity (*α*) for positive scores compared to negative scores, indicating that people tend to credit themselves for successes more readily than attribute failures to their actions. However, by including a task in which we could measure participants’ motor performance, our study expanded on this finding by showing that one of the main drives of the bias stems from distorted perception, independently of the positive reward score. Another study found that participants exhibited higher error sensitivity for negative outcomes due to self-attribution bias, as they needed good results to earn rewards and thus learned more from failures (Dorfman et al., 2019, 2021).

In contrast, our study required participants to achieve positive scores and estimate the task state accurately. Feedback was implicit, with scores providing only partial information on hit locations. A negative score could indicate either a no-error trial in the random condition or an error in the skill condition, and motor noise further disconnected outcomes from apparent success or failure. Consequently, participants focused more on transparent, informative trials. While Dorfman et al. emphasised learning from negative outcomes due to self-attribution, our study highlights how task structure and outcomes interact with self-attribution bias to shape error assignment and learning (Kim et al., 2019; Lee et al., 2024).

How does a self-attribution bias influence future choice? We found that misjudgements regarding perceptual abilities led participants to adjust their overall behavioural strategy. Specifically, they were more likely to change their choice if they believed the hidden state was skill-based but received a negative score or if they believed it was random and received a positive score. An effect that our model captured. These findings illustrate how erroneously attributing successes to personal skill and failures to external factors can significantly affect subsequent behavioural strategies, often resulting in future suboptimal decisions (Garrett & Daw, 2020). Our approach may thus provide a well-controlled framework to study self-attribution bias in the broader context of marketing, managerial and financial decision-making (Bossaerts, 2009; Karmarkar & Plassmann, 2019).

Using confidence ratings collected at the trial-by-trial level, we studied the relationship between self-attribution bias and metacognitive introspection. First, we confirmed that confidence, as predicted by classical signal-detection theory (SDT), was higher for correct inference decisions than for incorrect ones (Fleming, 2024; Maniscalco & Lau, 2012). Interestingly, we also detected an effect of valence on confidence – participants were more confident after positive feedback. While standard SDT does not predict these effects, they have been reported in the literature (Kaanders et al., 2022; Lebreton et al., 2019; Salem-Garcia et al., 2023). These results are consistent with a post-hoc positive bias, reflecting a positive post-decision evaluation of one’s actions.

Although we designed and fit the computational model only on inference accuracy, we found it could also capture key aspects of participants’ confidence judgments. Further simulations uncovered an important yet surprising underlying phenomenon (Supplementary Fig. 15). Participants’ confidence was closer to the model harbouring no perceptual distortion, implying a mechanistic dissociation between choice and confidence in self-attribution bias. This dissociation resonates with previous findings with obsessive-compulsive disorder patients, where individuals form accurate confidence judgements but fail to use this confidence to update action policies (Vaghi et al., 2017). From a different perspective, recent work found similar dissociations between choice accuracy and metacognitive evaluations that depended on the task context--simple perceptual decisions versus complex economic decisions (Lu et al., 2025).

Furthermore, after a hidden task state switch, confidence recovered faster in the skill-to-random transition (as opposed to the random-to-skill transition). This result was present even in the simulations with the oracle agent (using the true threshold, harbouring no biases in error sensitivity or constant term), suggesting that, in the context of our task, detecting an error was easier than confirming the absence of an error (Fu et al., 2023; Gehring et al., 1993; Kononowicz et al., 2022; Seidler et al., 2013). Finally, self-attribution bias affects confidence in a particular way, leading participants to be overconfident in attributing blame to external causes but not success to themselves.

In summary, our research reveals that self-attribution bias, a common yet subtle cognitive distortion, arises not only from processes of post-decision evaluation but, critically, also from perceptual distortions. An intriguing possibility is that perceptual distortions might be rooted in unequal or suboptimal evidence accumulation, an effect that eye-tracking and pupil-linked arousal processes could measure (Keung et al., 2019; Thomas et al., 2019). Our results offer fresh perspectives on how behavioural biases can emerge through multiple factors. They underscore the necessity of employing rigorous experimental designs alongside computational modelling to identify and understand these biases effectively. Such an approach will be crucial for developing targeted interventions to reduce or mitigate the impact of self-attribution bias.

## Methods

### Participants

We recruited 66 participants (23 female, mean ± SD age = 29.1 ± 8.8 years old, range 20-51 years old) to take part in this behaviour study. Participants were paid 3,000 JPY for a 1.5-hour participation. The ATR Institute International (Japan) Institutional Review Board approved the study protocol (ethics number #763). All participants provided written informed consent before beginning the experiments and were instructed they could withdraw their participation at any time. Participants were excluded from further analyses if they met any of the following exclusion criteria: (i) their accuracy about task state estimation in the main task was less than 54% --the minimum above-chance accuracy threshold given the total number of trials, determined with a binomial test; (ii) choosing the same option (‘skill’, or ‘random’) in more than 80% of the trials; (iii) reporting the same level of confidence in more than 80% in the trials. Thus, we selected 51 participants for the analyses reported in this paper (15 female, mean ± SD age = 28.2 ± 7.7 years old, range 20-46 years old).

### Task

The Whack-A-Mole arcade game inspired the main task: a player’s goal is to hit a mole while it briefly pops out from one hole out of a set of N holes on a board. We chose this game as it is a game of skill in which the player has to make rapid, precise movements to hit the mole, and it would lend itself well to outcome/score manipulations based on skill vs. luck (environment randomness).

Participants completed four task sessions: (1) mole hit and score prediction practice, (2) score prediction, (3) rule prediction practice, and (4) rule prediction main task. The first task was for participants to get used to hitting moles; the second was to evaluate the motor accuracy of participants and their ability to judge where they hit the mole and to calibrate the difficulty of the main task accordingly; the third was to let participants learn the structure of the main task, and the fourth was the main task. All tasks used a Windows PC with a touchscreen, and moles appeared and disappeared quickly, one by one, from one of the seven holes (see Figure 1). Each presentation started with a 100 ms period during which the mole moved up from the hole, followed by a 600 ms period during which the mole appeared in full, and finally, a 100 ms period during which the mole disappeared by descending in the hole. Participants were also told to use only the index finger of their right hand to touch the screen. Note that the period during which participants could hit a mole ranged from 200 ms from the start to the end (800 ms) of the mole presentation. If they missed a mole, the trial was aborted, but no penalty was applied, and another mole popped up at a different location after a delay of 200 ms.

#### Mole hit and score prediction practice

Participants were instructed to hit the mole as close to its centre as possible. When they hit a mole, concentric circles were displayed with a score of 0-100 points corresponding to the hit location. This practice session lasted 50 trials.

#### Score prediction

As in the initial practice, participants were instructed to hit the centre of the moles. After a hit, they predicted the points (0-100) with a slider without further clues (i.e., concentric circles were not shown). Participants also reported how confident they were about the correctness of their prediction on a Likert scale with four levels, with one being the lowest (guess) and four being the highest (certain). This prediction session lasted 100 trials. We used the hit location distribution to calculate the individualised threshold that would determine the main task’s binary score (positive/negative) in the skill condition. This threshold was calculated as the median hit location, i.e., the distance from the mole centre.

#### Rule prediction practice

Participants were instructed to hit the mole quickly before it disappeared. There were two task states: ‘skill’ and ‘random’. In the ‘skill’ state, the hit location directly determined the score. If participants hit closer to the centre than the threshold, they obtained a positive score, while they obtained a negative score if they hit further away but within the mole area. The threshold was individually calibrated for each participant from the preliminary score prediction task, such that ∼50% of mole hits would fall within the mole’s central area (hit location < threshold). Importantly, participants did not know the exact location of the threshold. In the ‘random’ state, scores were entirely random, i.e., there was an equal probability of getting a positive or negative score at any hit location. Participants had to estimate the current task state based on the displayed binary score (good/bad) and their belief about their hit location. During this practice session, they received feedback about the correctness of their answer at the end of each trial. This practice session lasted 80 trials.

#### Rule prediction main task

The main task had the same structure as the practice task described above, except that participants did not receive feedback about the task state (‘skill’ or ‘random’) or the correctness of their choices. Participants completed 560 trials; they were allowed to take a short break every 80 trials. The duration of a given sequence of trials within one hidden task state, i.e., between two state change points, followed a Gamma distribution with parameters k = 22.5, θ = 1.33, with a theoretical mean duration of 29.9 trials (true empirical mean across participants was 28.8 ± 1.5 trials). Based on performance, an extra reward was given to incentivise participants to hit the centre of the mole while correctly estimating the current task. Participants were told at the beginning of the task that they would receive a reward point each time they correctly estimated the task state and obtained a positive score. However, they were not informed about whether they were rewarded and how much until the end of the entire experiment. In the ‘random’ state, the reward rate was calculated as 50% of the correctly estimated trials (instead of considering actual empirical scores). The real calculation was not revealed to participants to prevent probabilistic biases. The maximum extra reward was 3,000 JPY, and participants received an average of 2,122 ± 296 JPY.

### Model-free analyses

#### General Behaviour

Inference accuracy in the task was calculated as the ratio of correct answers (‘skill’, ‘random’) given the true hidden task state (‘skill’, ‘random’). While the theoretical chance level was 0.5, we used a binomial test to determine at the individual level if a participant’s data was overall above chance or not. Given 560 total trials, with a probability 0.5 of being correct on any given trial, the above-chance inference accuracy threshold was thus 0.54. As mentioned in the exclusion criteria, any participant whose accuracy was lower than this threshold was removed from further analyses. As representative examples, we selected two random participants and plotted the time series of their choices given the hidden rule. The time series of participants’ choices was calculated as the moving average (backward window size = 15 trials) of the proportion of ‘skill’ choices. Overall, participants were similarly accurate in detecting skill and random states (see Supplementary Table 1).

#### Contextual biases in choice behaviour

To investigate whether participants displayed any choice bias based on the scores received, we plotted the ratio of ‘skill’ choices, accuracy, and confidence separately for each score. Confidence ratings were z-scored within each participant. Wilcoxon signed-rank test with continuity correction was used to test the difference between the two score conditions. We used repeated measures two-way ANOVA to test the effect of scores and task states and their interaction on inference accuracy or confidence separately. Wilcoxon signed-rank test with continuity correction was used to test the simple main effect of each factor when the interaction was significant. Because there were four pairwise comparisons (given two scores and two task states), p-values were adjusted with false-discovery rate (FDR) correction.

#### Task state change points and participants’ behavioural adaptation

To see how participants adapted their choice behaviour and confidence ratings to task state switches, we first aligned the data to the task change points. Next, we labelled the change points based on the direction of the switch, i.e., ‘skill’ → ‘random’ and ‘random’ → ‘skill’. We extracted the four trials preceding the change point for each switch direction and the eight trials following the change point. We averaged trials at each time point across occurrences within participants. The twelve-trial time courses within participants were then used across participants to compute the population mean and standard error of the mean for each switch direction. To statistically analyse how the switch direction affected inference accuracy and confidence following a task state switch, we used repeated measures two-way ANOVA with factors time (trials from the switch) and switch direction, as well as their interaction. We further tested the difference in accuracy or confidence at each trial after the switch using a Wilcoxon signed-rank test with continuity correction, adjusted with FDR for multiple comparisons correction across the eight trials.

To investigate the underlying factors that caused participants to switch their choices, we plotted the subjective switching probability based on the previous choice (‘skill’, ‘random’) or confidence (‘low’, ‘high’) and the score (‘positive’, ‘negative’). Note we binarised confidence into high and low levels through a median split within each participant. The effect on switch probability of the two factors, choice/confidence and score, and their interaction, was tested using repeated measures two-way ANOVA. We applied Wilcoxon signed-rank test with continuity correction and FDR adjustment to each pair if the interaction was significant.

#### Hitting behaviour

We investigated participants’ mole-hitting patterns. This analysis examined whether participants might have changed where they hit the mole (consciously or unconsciously) depending on the score and inferred task state. To this end, for each trial, we extracted the distance between the hit location and the centre of the mole. We normalised the obtained value by the within-participant median distance. Using repeated measures two-way ANOVA, we tested the difference in the distance between the current and next trial. If the interaction was significant, we applied a Wilcoxon signed-rank test with continuity correction and FDR adjustment for multiple comparisons to each pair.

### Model-based Analyses

We developed a leaky error-accumulation model with separate error sensitivity and constant terms for each outcome type [positive vs. negative (den Ouden et al., 2013; Lefebvre et al., 2017)]. Here, we defined prediction errors following the intuition that participants have an internal estimate of how well they can hit the mole’s centre, which depends on a subjective threshold parameter. This estimate helps them make an internal decision on whether they were more likely to have hit the centre or the edge of the mole.

#### Full model

The model features a leaky decision evidence accumulator, updated on each trial based on an error signal resulting from the mismatch between the binarised hit location (centre, edge) and the score (positive, negative). To judge if the hit location was in the centre, the model assumes participants have an internal, subjective estimate of the size of the central area, i.e., a threshold parameter that reflects the hypothetical separation between the central and the outer regions. Accordingly, there will be no error if participants hit the centre and obtain a positive score or hit outside the central area and get a negative score. Other cases (such as hitting the centre and obtaining a negative score) will result in an error signal. In this context, the model is closest to the ground truth when the threshold parameter is identical to the true threshold used in the task to determine scores in the skill state. Thus, participants will be sub-optimal if they overestimate (subjective threshold > true threshold) or underestimate (subjective threshold < true threshold) how well they hit the centre. The model evaluates if the agent hits the centre with the equation:

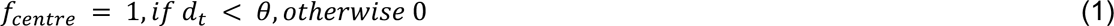

where *t* represents the trial, *d* is the distance between the hit location and the centre of the mole, and *θ* is the free parameter representing the subjective threshold.

The error *e_t_* is calculated using the centre-edge estimate and the score *o_t_* obtained at trial *t*:

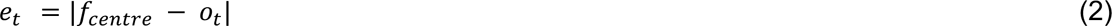

The model accumulates the error through a hidden variable *z_t_*, and outputs the choice probability *P*:

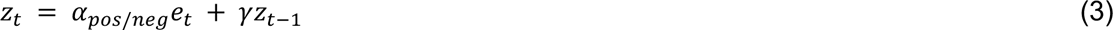

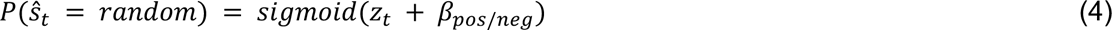

where 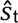 is the choice, and (*α*_pos_, *α*_neg_, *β*_pos_, *β*_neg_, *γ*) are free parameters; *α* represents the error sensitivity for the current error observation (differently for positive and negative scores), *β* is a constant term modulating the decision boundary on the inference choice (differently for positive and negative scores), and *γ* is the retention factor. Thus, the full model has six free parameters in total. Note that, for simplicity, we did not explicitly include *d_t_* in the model’s computation of choice probability, because *d_t_* would produce an additional non-linearity (the threshold being elsewhere than halfway between centre and edge) which would require additional parameters.

Finally, we calculate negative entropy as:

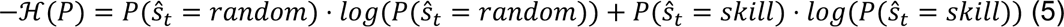

We used negative entropy z-scored within participants as a model measure of confidence to compare with participants’ actual confidence ratings. However, it is important to note here that confidence was not explicitly modelled (i.e., it was not part of the model fitting procedure) and remained a simple additional read-out.

#### Parameter estimation

We fit the model separately for each participant. The model takes the score sequence as input, and the trial-wise hit location is expressed as the distance from the centre, which outputs the choice probability. Consequently, we computed the cumulative negative log-likelihood over all time points within participants. For each participant, we estimated all free parameters (*α*_pos_, *α*_neg_, *β*_pos_, *β*_neg_, *γ*, and *θ*) simultaneously by minimising the negative log-likelihood with a numerical minimisation method. The minimisation was done with the *constrOptim* function of R using the Nelder-Mead algorithm. Furthermore, the error sensitivity *α* and constants *β* were constrained within the range [-20, 20], the retention factor *γ* within the range [0, 1], and the threshold *θ* within the range [0, 43] (note that 43 represents the smallest integer superior to the farthest hit location across all participants). We fit the parameters with ten different sets of random initialisations and used the best result as the final initialisation to fine-tune the parameters and perform model comparisons.

#### Alternative models and model comparisons

We compared the full model with a range of alternative, simpler models. These were created by simplifying the main full model, such as removing specific parameters or the positive/negative asymmetry. The full list of models is displayed in Supplementary Table 2.

All models were fitted to each participant’s data using the same procedure described above. The randomly initialised parameters were fitted using the Nelder-Mead algorithm with a constrained range and fine-tuned using the best ones as starting points. We calculated the Akaike Information Criterion (AIC) and Bayesian Information Criterion (BIC) for each participant and model and used them to evaluate goodness of fit. We used a Wilcoxon signed-rank test to test the AIC difference between each model and the full model (Supplementary Fig. 5).

#### Parameter recovery

We performed a parameter recovery analysis to validate the model’s reliability. First, we defined 100 sets of ground truth parameters; each set was randomly generated by sampling each parameter from a normal distribution whose mean and standard deviation were calculated from participants’ estimated parameter values. Next, we generated a sequence of simulated choices for each parameter set, using the original sequence of hit locations and scores as input to the model. As a result, we obtained 5100 simulated sequences (51 original sequences X 100 simulations each). The model fitting procedure was applied to each simulated sequence, thus obtaining pairs of ground truth and estimated parameters. Finally, we calculated Spearman’s rank correlation coefficient for each parameter and generated the corresponding confusion matrix (Supplementary Figure 7). Results indicate good recovery performance, with correlations in the range of *r* = 0.74, 0.88. Instead, correlations between pairs of unrelated parameters resulted in values near zero, in the range *r* = [-0.06, 0.09].

#### Model simulations

We performed additional model simulations with arbitrary parameter settings to validate the model findings on key behavioural indicators (Palminteri, Wyart, et al., 2017). For all these simulations, the model sequentially computed *p_t_*, the probability of choosing “random” for each trial based on a given parameters set, using the participants’ scores and hit locations (distance from the mole centre) as inputs. As a result, the model outputs a time series of probabilities for each participant. Additionally, trial-by-trial entropy was calculated using equation (5). To prevent issues with logarithm calculations when *p_t_* = 1, *p_t_* was clipped to a maximum of 1 − 10^−5^. Accuracy for each trial was defined as *p_t_* when the true state was random and 1 − *p_t_* when the true state was skill. The negative entropy values were normalised within participants and used as a proxy for confidence.

Our objectives were to examine the impact of the asymmetry in parameter estimates for error sensitivity and constant terms and of threshold overestimation on performance. We thus focused on two sets of parameters: the threshold (*θ*) and the error sensitivity/constant term parameters (*α* and *β*).

To this end, we implemented two conditions for the threshold modifications to assess overconfidence in hitting ability. The Baseline (Overestimated Threshold) Condition used a *θ* value estimated through model fitting based on the observed behaviour, which systematically overestimates the true threshold. In the True Threshold Condition, the *θ* parameter was set to each participant’s true threshold, as determined by the task specifications, providing a benchmark that reflects optimal or unbiased performance. To evaluate the effect of asymmetrical error sensitivity and constant terms, we introduced two additional conditions alongside a baseline for comparison, which uses the estimated *α* and *β* from the model fitting. In the Mean Parameters Condition, we removed the asymmetry by averaging the parameter estimates; expressly, *α* was set to the mean of the positive and negative *α* values, and *β* was set to the mean of the corresponding *β* values.

Based on these conditions, we conducted eight simulations as follows: (i) with subjective threshold, score-dependent error sensitivity and constant term; (ii) with true threshold, score-dependent error sensitivity and constant term; (iii) with subjective threshold, score-dependent error sensitivity and single constant term; (iv) with true threshold, score-dependent error sensitivity and single constant term; (v) with subjective threshold, single error sensitivity and score-dependent constant term; (vi) with true threshold, single error sensitivity and score-dependent constant term; (vii) with subjective threshold, single error sensitivity and constant term; (viii) with true threshold, single error sensitivity and constant term.

## Data and code availability

Behaviour data and code used for the analyses are available at https://github.com/ATR-decnef/WAM_publish

## Acknowledgements

We thank Peter Dayan for comments on an earlier draft, Nizar Mathli and Shihab Ahmed for help with task preparation, and Kaori Nakamura for help with participants’ recruitment.

## Author contributions

M.T., B.D.M. and A.C. conceived the study; N.O. collected data and performed data analysis with supervision from B.D.M. and A.C.; T.K. and S.I. provided critical feedback on data analysis and computational modelling; N.O., B.D.M. and A.C. drafted and wrote the manuscript with input from T.K. and S.I.; A.C. acquired funding; B.D.M. and A.C. supervised the project.

