## Supplementary materials (tables and figures) for "Blaming Luck, Claiming Skill: Self-Attribution Bias in Error Assignment"

| | | task state | |
|---|---|---|---|
| | | skill | random |
| choice | skill | 0.70 ± 0.13 | 0.35 ± 0.13 |
| | random | 0.30 ± 0.13 | 0.65 ± 0.13 |

**Supplementary Table 1**. Confusion matrix of task state and choices. Each cell is the ratio of choices within the relevant task state. The ratios within each task state sum to one within participants and are reported as mean ± STD computed across participants. Wilcoxon signed rank test on diagonal elements, i.e., true positive and true negative rates: Z = 1.87, P = 0.061.

| Model | Free parameters | Mean AIC / BIC / LL |
|---|---|---|
| **Full model** | $\boldsymbol{\alpha_{pos}, \alpha_{neg}, \beta_{pos}, \beta_{neg}, \gamma, \theta}$ | **463.9 / 489.9 / -226.0** |
| Single constant term | $\alpha_{pos}$, $\alpha_{neg}$, $\beta$, $\gamma$, $\theta$ | 483.3 / 504.9 / -236.7 |
| Single error sensitivity | $\alpha$, $\beta_{pos}$, $\beta_{neg}$, $\gamma$, $\theta$ | 502.0 / 523.7 / -246.0 |
| Single error sensitivity, constant | $\alpha$, $\beta$, $\gamma$, $\theta$ | 519.3 / 536.6 / -255.7 |
| Fixed, true threshold | $\alpha_{pos}$, $\alpha_{neg}$, $\beta_{pos}$, $\beta_{neg}$, $\gamma$ | 525.5 / 547.1 / -257.7 |
| No retention factor | $\alpha_{pos}$, $\alpha_{neg}$, $\beta_{pos}$, $\beta_{neg}$, $\theta$ | 574.3 / 595.9 / -282.1 |

**Supplementary Table 2**. List of models used in the computational analysis of behaviour. The full model, highlighted in bold, provided the best fit across participants (lowest AIC, BIC, LL).

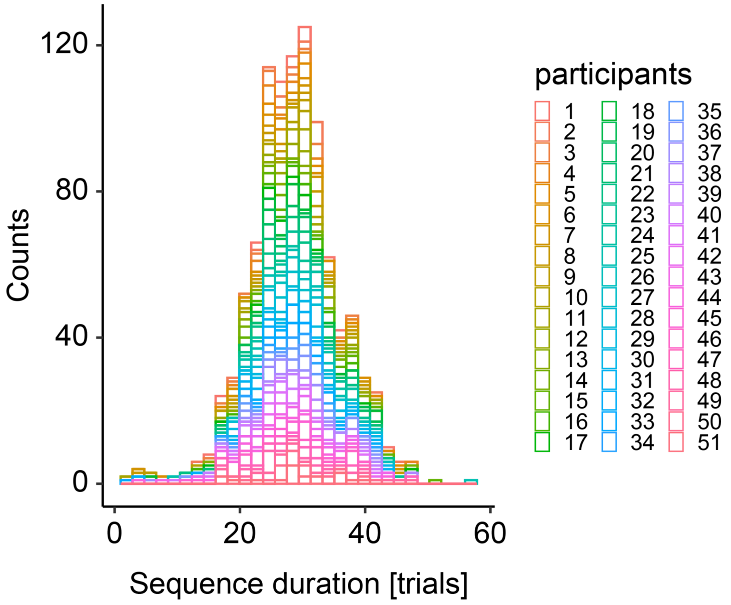


**Supplementary Figure 1.** Histogram of durations in trials of hidden task state sequences. The mean and standard deviation of individual durations was 28.8 ± 1.5. Each colour represents one participant (N = 51).

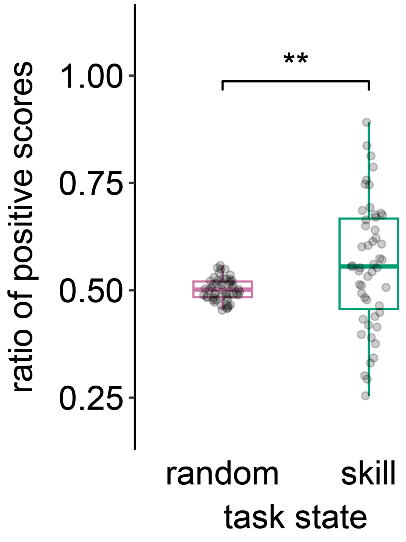


**Supplementary Figure 2.** The task was calibrated such that participants would obtain roughly 50% positive scores in both skill and random states. Note that while there was individual variability, overall, at the group level the ratio of positive scores was relatively similar across states (Wilcoxon signed-rank test, Z = 2.58, p = 0.01).

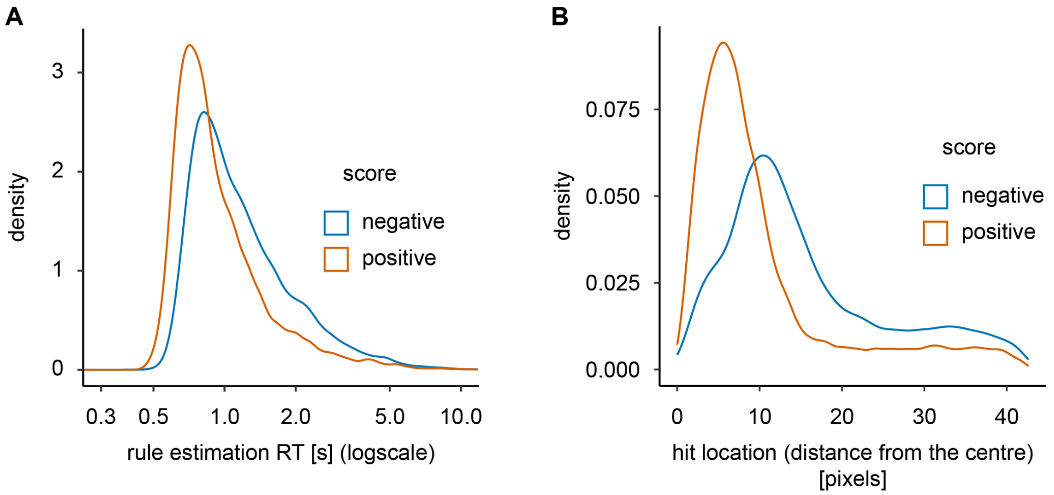


**Supplementary Figure 3.** (A) Distribution of log-transformed reaction times in rule estimation, divided by score (positive, negative). (B) Distribution of hit locations, expressed in pixels as the distance from the mole centre and divided by score (positive, negative).

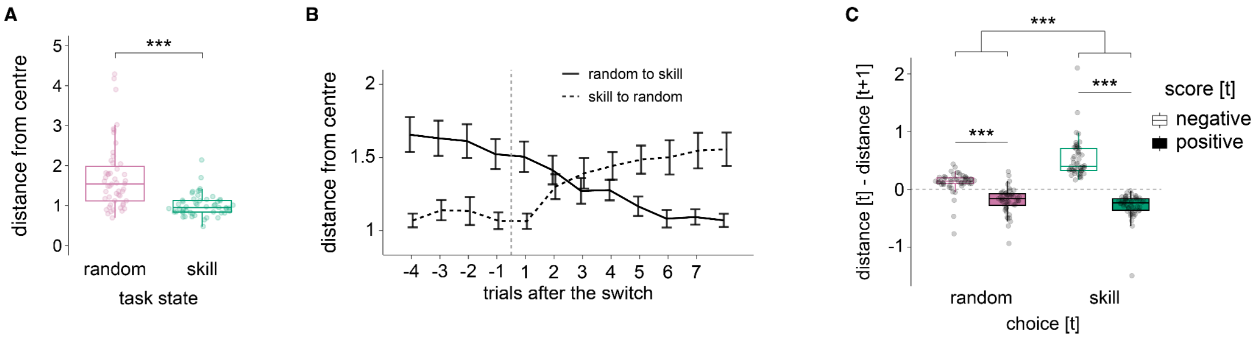


**Supplementary Figure 4.** Participants' hit location patterns expressed as the distance of the hit from the mole centre. (A) distance from the centre divided by hidden task state (random, skill). (B) distance from the centre around task state switches, plotted from -4 trials before the switch to +8 trials after the switch. The trajectories are plotted separately for the two types of transition, from random to skill and skill to random. (C) cross-trials dynamics of participants' hit behaviour, separately for trials in which participants chose random or skill after obtaining a negative or positive score. The plot shows that participants adjusted their hit locations accordingly. After receiving a negative score, participants tended to hit closer to the centre (increased precision of hits). In contrast, after receiving a positive score, participants tended to hit further away from the centre (a relaxation of the precision).

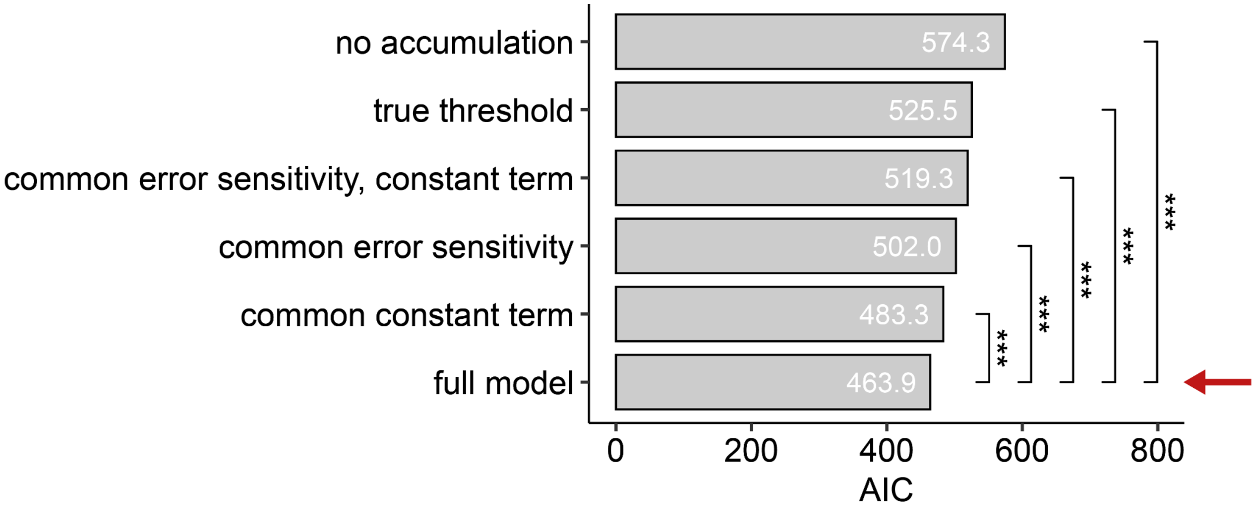


**Supplementary Figure 5.** Comparison of the full model against alternative models, in which specific parameters, or sets thereof, were fixed (e.g., in which the error sensitivity $\alpha$ was constrained to be score-independent). We used the Akaike Information Criterion (AIC) (Akaike 1974; Symonds and Moussalli 2010) to compare models. Note that lower AIC values mean better model fit. The plot shows each model's average AIC values (computed across participants). Wilcoxon signed-rank test, with FDR correction for multiple comparisons, demonstrated that the full model provided a significantly better fit for participants' data. It also showed that the subjective threshold parameter played a more important role than the constant term or the error sensitivity (both reflecting a positivity bias) due to the larger difference in AIC compared to the full model. The red arrow indicated the best-fit model, *** $P < 0.001$.

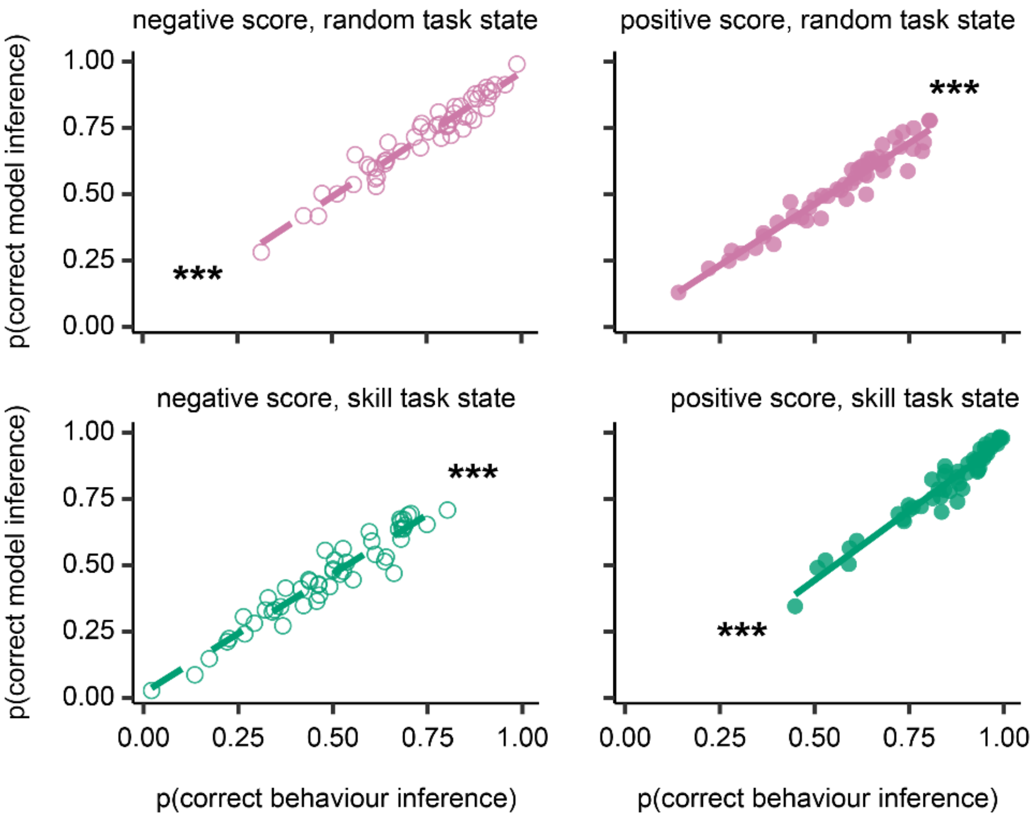


**Supplementary Figure 6.** Quantitative correspondence between participants' behaviour and the model's simulated behaviour in terms of *p*(correct state inference). Each subplot represents one of the four conditions based on scores (negative, positive) and hidden task states ('random', 'skill'). From top left to bottom right: negative score, random task state (r = 0.96, $P < 10^{-10}$); positive score, random task state (r = 0.96, $P < 10^{-10}$); negative score, skill task state (r = 0.96, $P < 10^{-10}$); positive score, skill task state (r = 0.96, $P < 10^{-10}$). Each plot N = 51, circles represent individual participants, and solid/dotted lines represent the linear fit.

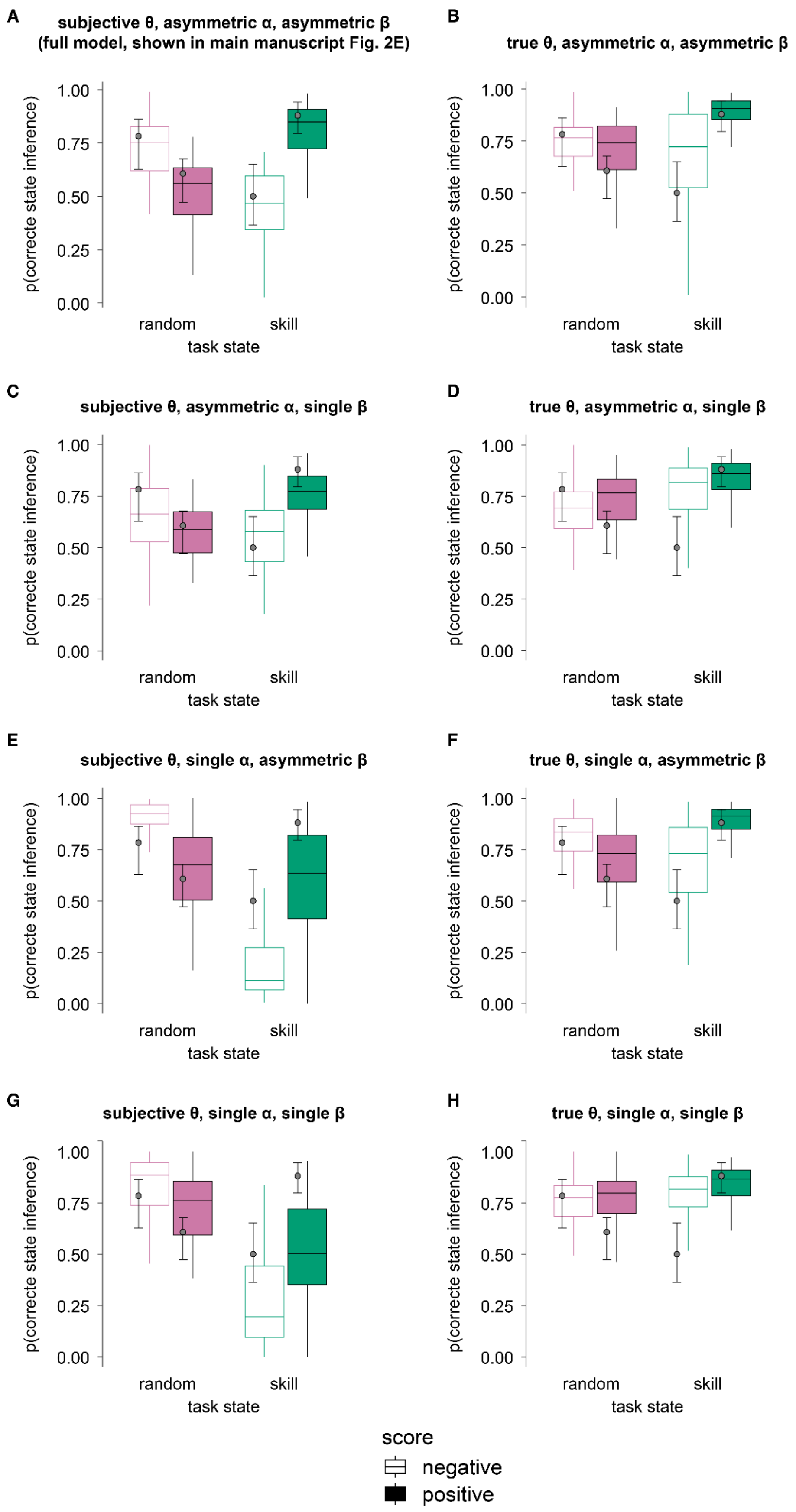
A
subjective θ, asymmetric α, asymmetric β
(full model, shown in main manuscript Fig. 2E)
B
true θ, asymmetric α, asymmetric β
C
subjective θ, asymmetric α, single β
D
true θ, asymmetric α, single β
E
subjective θ, single α, asymmetric β
F
true θ, single α, asymmetric β
G
subjective θ, single α, single β
H
true θ, single α, single β
p(correcte state inference)
1.00
0.75
0.50
0.25
0.00
random
skill
task state
score
negative
positive

**Supplementary Figure 7.** Inference accuracy, plotted by hidden task state and score. Simulations with specific parameter settings to evaluate the effect of cognitive strategies on inference accuracy. We performed eight simulations based on three parameters, with two possible settings each. Across plots, the small circles represent participants' average inference accuracy, and the error bars are the standard error of the mean. The boxplots represent the simulation median and interquartile ranges. (A) Simulating the best fitting model with subjective threshold, asymmetric (score-dependent) error sensitivity, and asymmetric (score-dependent) constant term. This simulation is the same as reported in the main manuscript in Fig 2E. Note the close mapping between participants' behaviour and simulation results. (A, C, E, G) Simulations based on the model with the subjective threshold, modifying the error sensitivity and/or the constant term. (B, D, F, H) Simulations based on the model with the true threshold, modifying the error sensitivity and/or constant term. (H) Oracle agents use the true threshold and equal weights across positive and negative scores for error sensitivity and constant term. For all panels, simulation results are plotted as boxplots, with the box representing the median, first and third quartiles and whiskers representing the minimum and maximum of the data range, while participants results are plotted as circles representing the median and error bars representing the first and third quartiles.

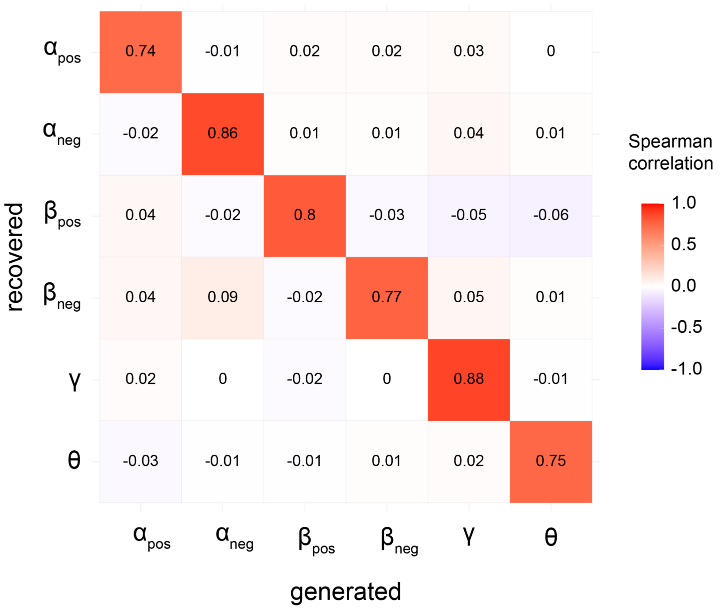


**Supplementary Figure 8.** Parameter recovery analysis, showing the confusion matrix with correlations between original (generated) parameters and recovered parameters.

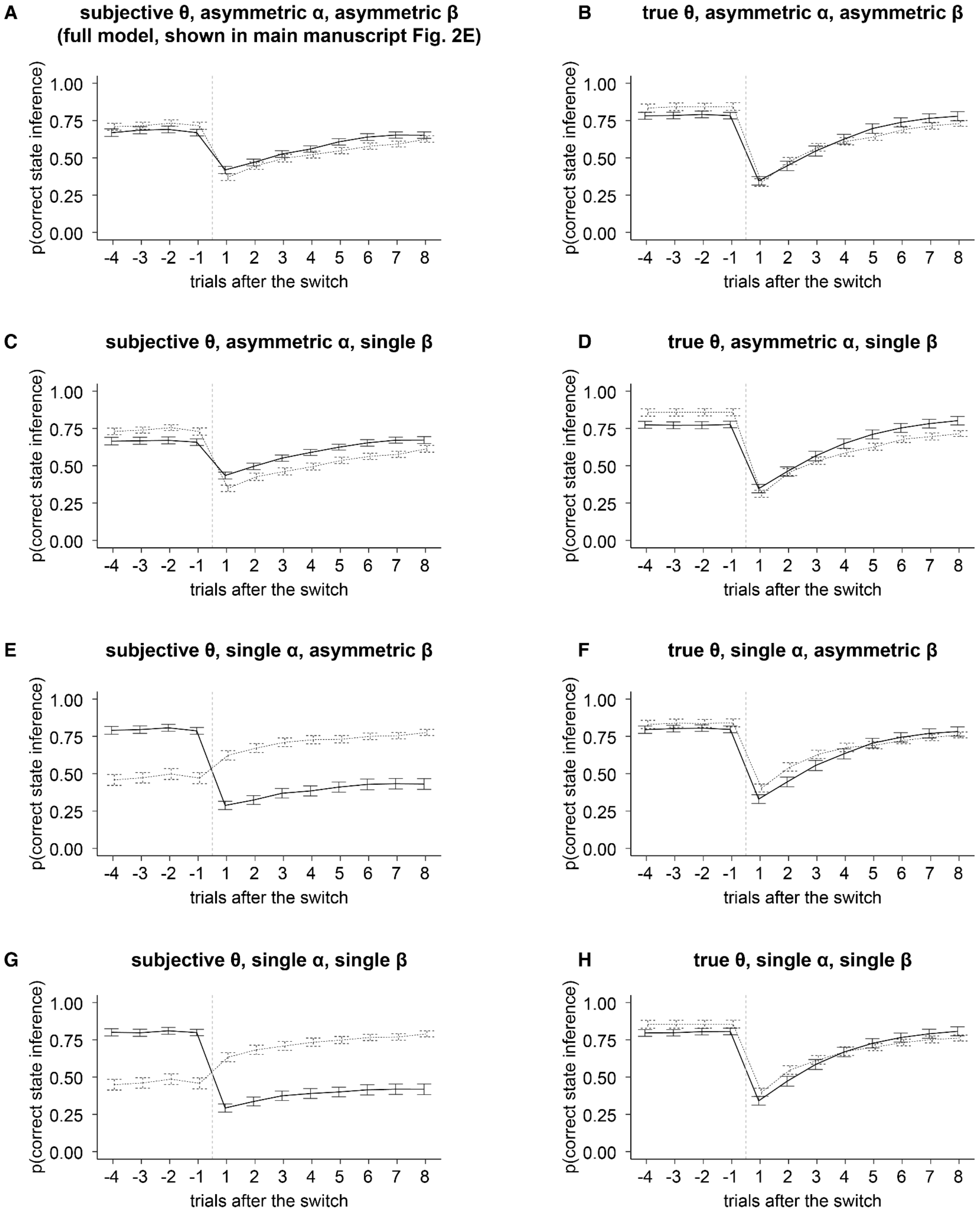


**Supplementary Figure 9.** Inference accuracy, plotted around hidden task state switches. Simulations with specific parameter settings to evaluate the effect of cognitive strategies on inference accuracy. We performed eight simulations based on three parameters, with two possible settings each. Across plots, the small circles represent participants' average inference accuracy, and the error bars represent the standard error of the mean. The boxplots represent the simulation median and interquartile ranges. (A) Simulating the best fitting model with subjective threshold, asymmetric (score-dependent) error sensitivity, and asymmetric (score-dependent) constant term. This simulation is the same as reported in the main manuscript in Fig 2E. Note the close mapping between participants' behaviour and simulation results. (A, C, E, G) Simulations based on the model with the subjective threshold, modifying the error sensitivity and/or the constant term. (B, D, F, H) Simulations based on the model with the true threshold, modifying the error sensitivity and/or constant term. (H) Oracle agents use the true threshold and equal weights across positive and negative scores for error sensitivity and constant term.

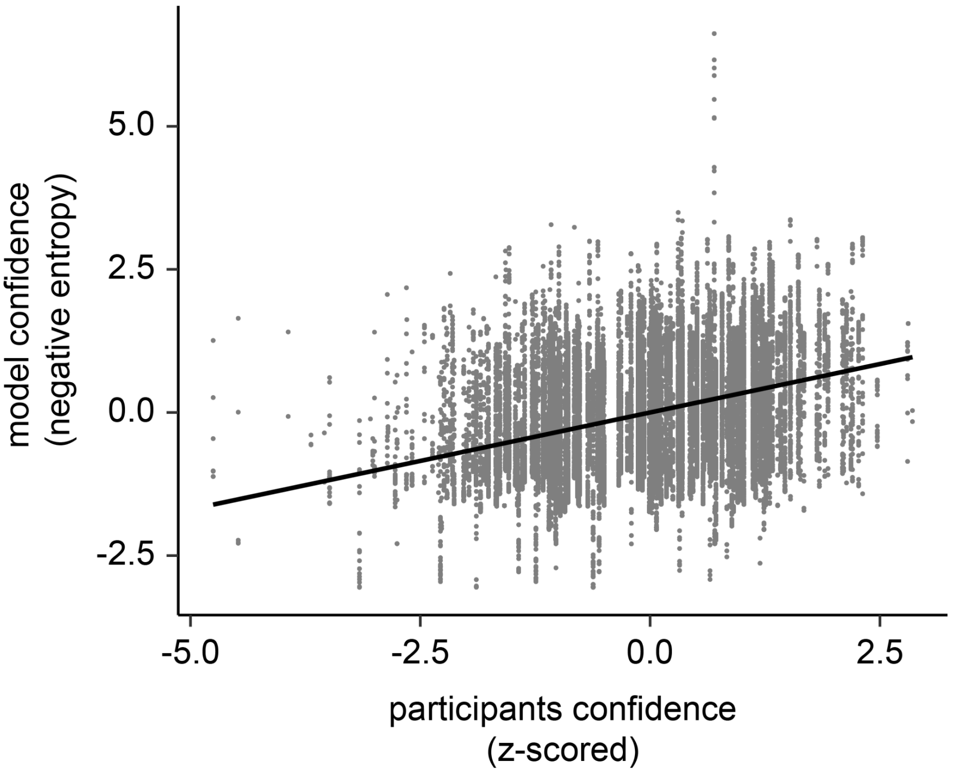


**Supplementary Figure 10.** Correspondence between participants' confidence judgements and negative entropy of the model's decision output. Importantly, the model was not optimised based on confidence but solely based on the inference choices. The model's confidence was taken simply as the negative entropy of the decision output (since entropy signals the uncertainty in that decision). We evaluated the linear relationship with a linear mixed-effect model [Wilkinson formula *negative_entropy ~ confidence + (confidence | subjID)*]. The factor confidence was significant (estimate = 0.34, std = 0.026, $t_{50}$ = 12.86, $P$ < 0.001).

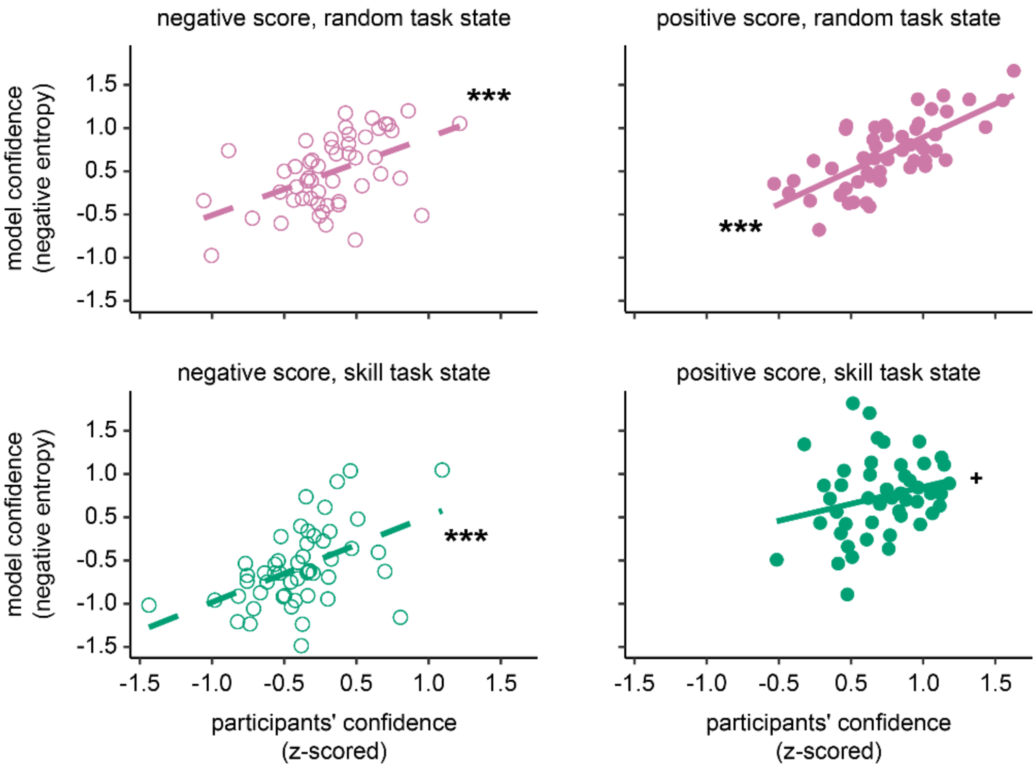


**Supplementary Figure 11.** Quantitative correspondence between participants' behaviour and the model's simulated behaviour in terms of p(correct state inference). Each subplot represents one of the four conditions based on scores (negative, positive) and hidden task states ('random', 'skill'). From top left to bottom right: negative score, random task state ($r = 0.49$, $P < 0.001$); positive score, random task state ($r = 0.69$, $P < 10^{-4}$); negative score, skill task state ($r = 0.52$, $P < 0.001$); positive score, skill task state ($r = 0.26$, $P = 0.069$). Each plot N = 51, circles represent individual participants, and solid/dotted lines represent the linear fit.

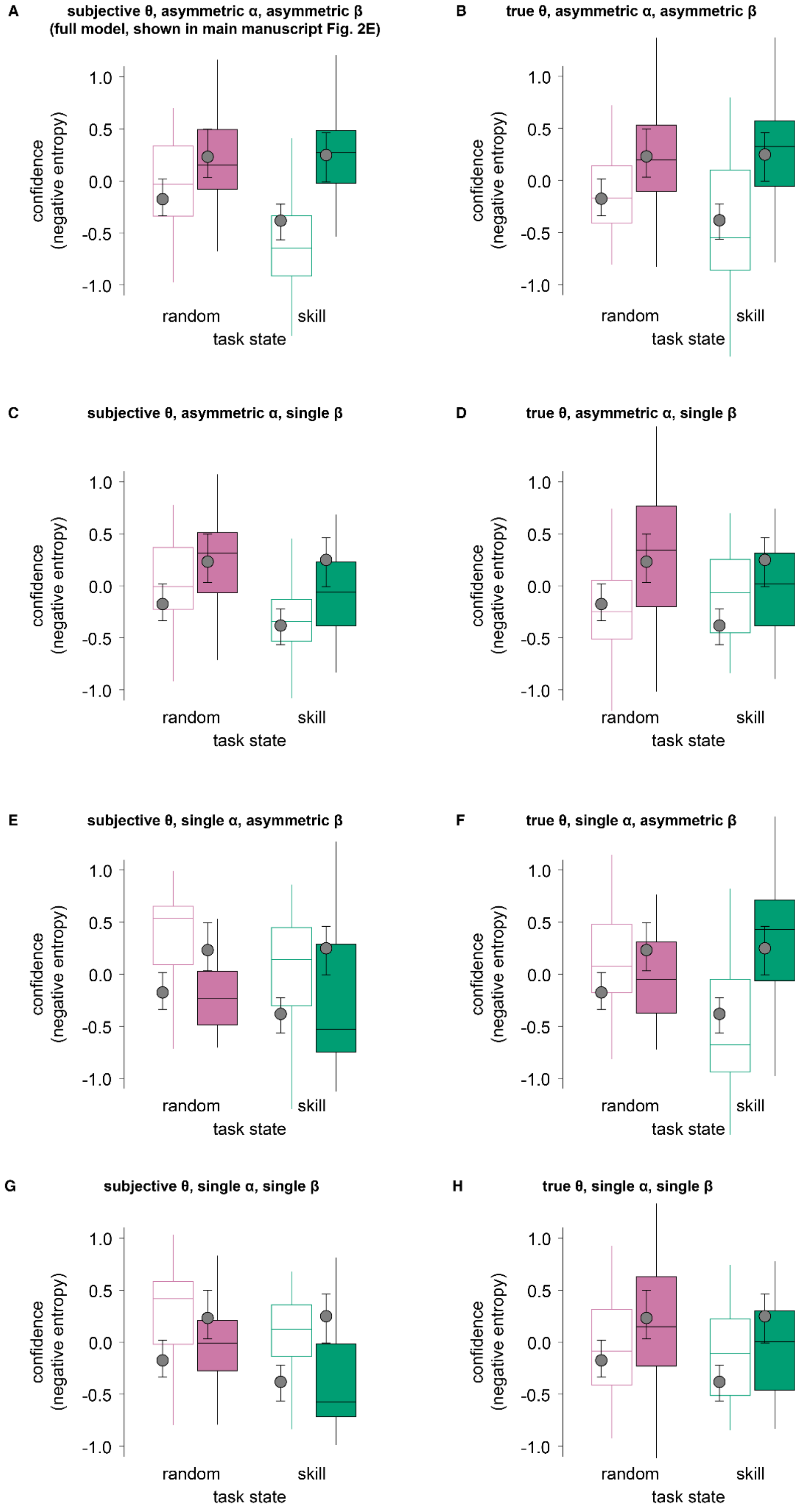
A
subjective θ, asymmetric α, asymmetric β
(full model, shown in main manuscript Fig. 2E)
B
true θ, asymmetric α, asymmetric β
C
subjective θ, asymmetric α, single β
D
true θ, asymmetric α, single β
E
subjective θ, single α, asymmetric β
F
true θ, single α, asymmetric β
G
subjective θ, single α, single β
H
true θ, single α, single β
confidence
(negative entropy)
1.0
0.5
0.0
-0.5
-1.0
random
skill
task state

**Supplementary Figure 12.** Confidence (negative entropy), plotted by hidden task state and score. Simulations with specific parameter settings to evaluate the effect of cognitive strategies on confidence. (A) Simulating oracle agents, using the true threshold and equal weights across positive and negative scores for error sensitivity and bias. Confidence shows minimal variation across hidden task states and scores. (B) Simulating agents with the true threshold but with a score-dependent error sensitivity and bias. Confidence again displays the original effect of scores, with higher confidence for positive scores than negative ones. (C) Simulating agents using the subjective threshold and score-independent error sensitivity and bias. Confidence shows the opposite pattern, being lower for positive scores and higher for negative scores and overall lower in skill compared to random task state. (D) Simulating agents using the true threshold and inverted error sensitivity and biases (instead of the original weights being larger for positive than negative scores, weights are now larger for negative than positive scores). In this case, the confidence pattern reverses entirely compared to the original result, with higher confidence for negative than positive scores, regardless of task state. For all panels, simulation results are plotted as boxplots, with the box representing the median, first and third quartiles and whiskers representing the minimum and maximum of the data range, while participants results are plotted as circles representing the median and error bars representing the first and third quartiles.

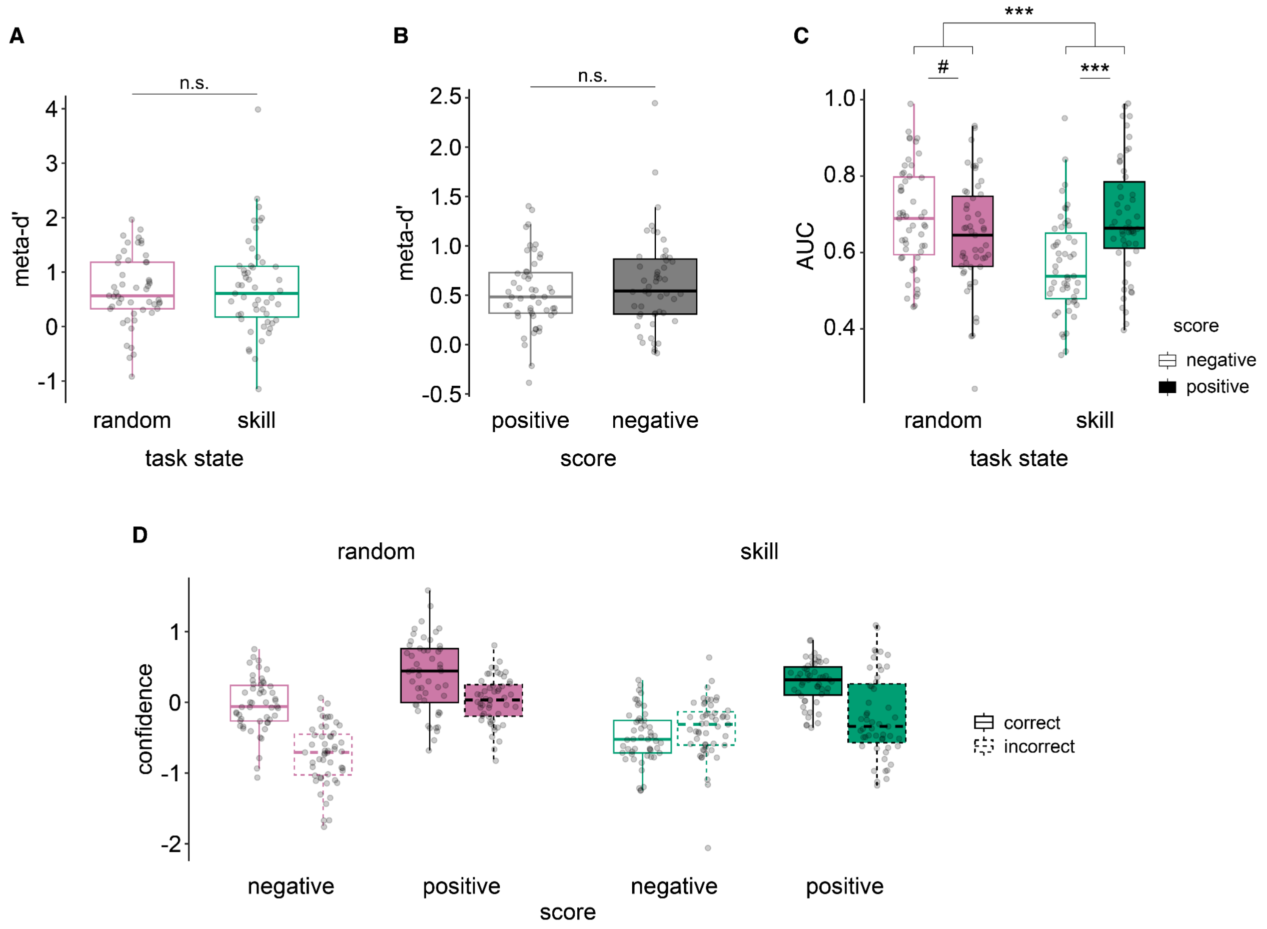


**Supplementary Figure 13.** Metacognition. (A) Formal meta-d' analysis for each task state (random and skill). Participants had equal metacognitive capacity across states. HMeta-d was used to calculate meta-d' (Fleming 2017). (B) Formal meta-d' analysis based on the score obtained (negative, positive). Participants had equal metacognitive capacity across score types. (C) Area under the curve (AUC) analysis separately for each score-task state combination. Note that because we are now looking at each condition separately, it is no longer possible to compute meta-d' (which requires data from both responses within each condition of interest). We thus computed AUC on the confidence-choice response operating curve. Note the steep drop in AUC (metacognitive ability) specific to the negative scores in the skill task state. (D) Showing the underlying plot (C) data, notice the absence of a difference in confidence between correct and error trials, specifically after the negative scores in the skill task state. For all boxplots, the box represents the median, first and third quartiles, and whiskers represent the minimum and maximum of the data range; scatter plots represent participants' individual data points.

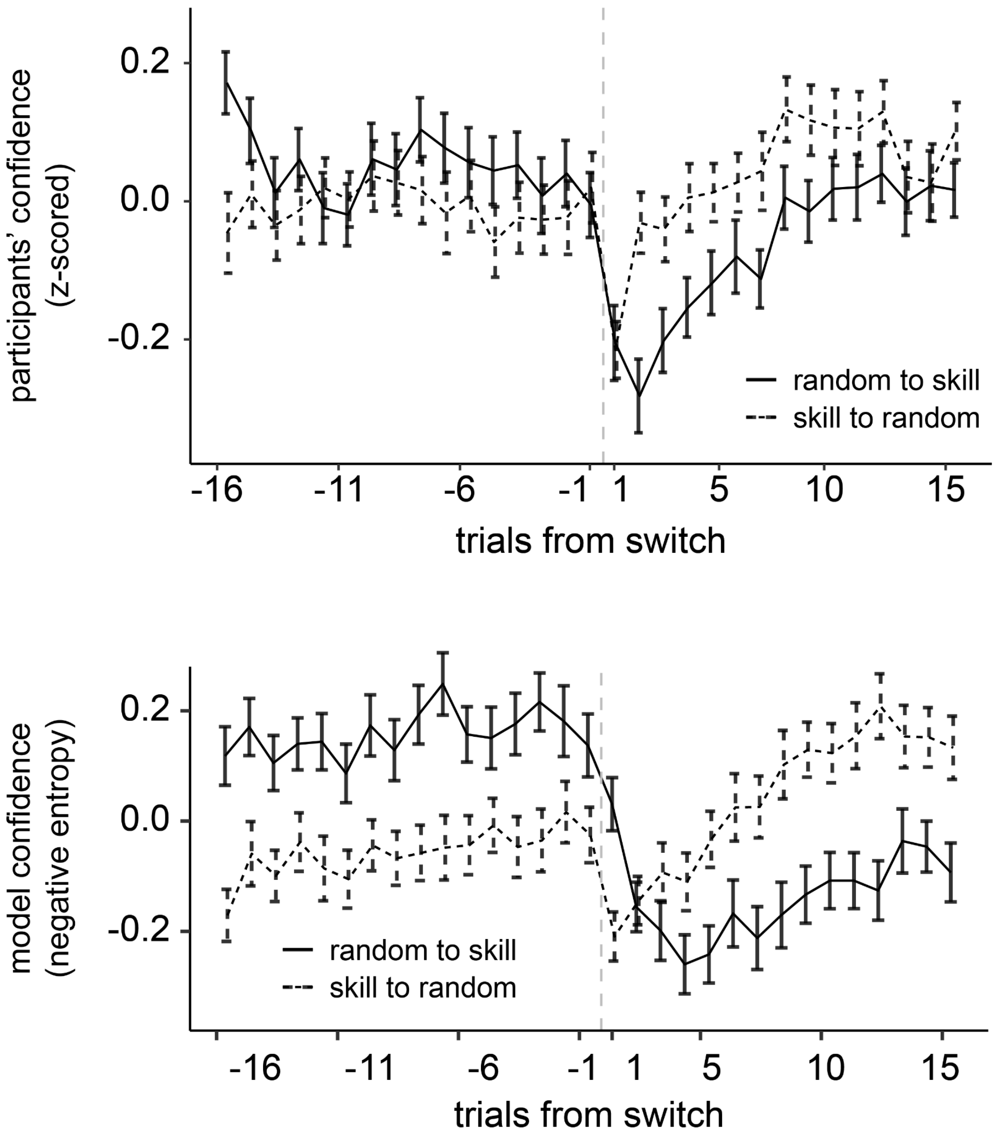


**Supplementary Figure 14.** Time series of participants' confidence judgements (top) and model confidence (bottom) around hidden task state switches (e.g., random → skill or skill → random). The central line represents the mean across participants, and the error bars represent the standard error of the mean. Confidence was first z-scored and averaged within each participant/simulation. In both cases, the random condition leads to higher average confidence than the skill condition. While this effect was stronger in the model results, it was also clearly present in participants' data.

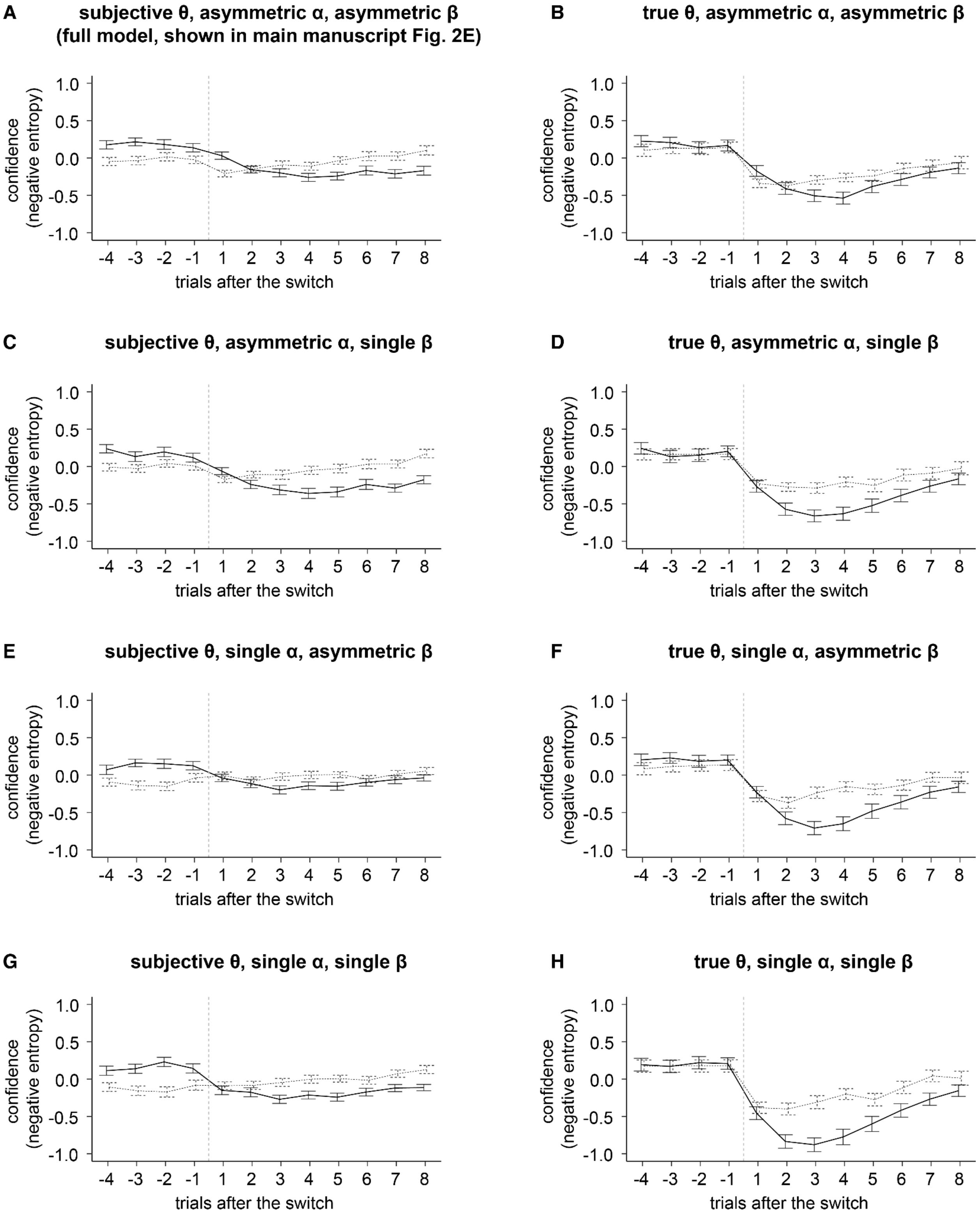


**Supplementary Figure 15.** Confidence (negative entropy), plotted around hidden task state switches. Simulations with specific parameter settings to evaluate the effect of cognitive strategies on inference accuracy. We performed eight simulations based on three parameters, with two possible settings each. Across plots, the small circles represent participants' average inference accuracy, and the error bars are the standard error of the mean. The boxplots represent the simulation median and interquartile ranges. (A) Simulating the best fitting model with subjective threshold, asymmetric (score-dependent) error sensitivity, and asymmetric (score-dependent) constant term. This simulation is the same as reported in the main manuscript in Fig 2E. Note the close mapping between participants' behaviour and simulation results. (A, C, E, G) Simulations based on the model with subjective threshold, modifying the error sensitivity and/or the constant term. (B, D, F, H) Simulations based on the model with the true threshold, modifying the error sensitivity and/or constant term. (H) Oracle agents use the true threshold and equal weights across positive and negative scores for error sensitivity and constant term.

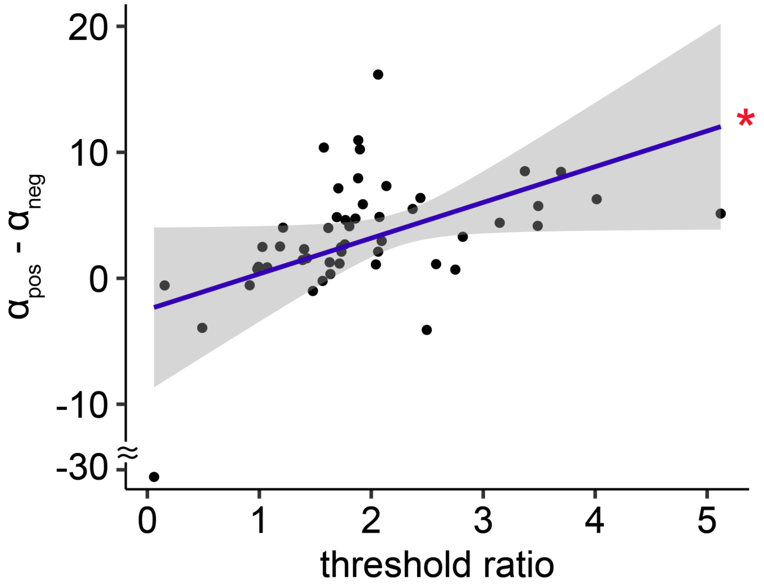


**Supplementary Figure 16.** The plot shows the linear relationship between the perceptual distortion (represented as the ratio between the subjective and true thresholds) and the strength of the positivity/confirmation bias (defined as the difference between the error sensitivity for positive vs negative outcomes). We used robust regression to evaluate the strength of the relationship (estimate = 2.84 ± 1.40, $t_{49}$ = 2.02, $P$ = 0.049). Dots represent individual participants, the line the linear fit, and the red asterisk $P$ < 0.05.
